# Comparison and evaluation of statistical error models for scRNA-seq

**DOI:** 10.1101/2021.07.07.451498

**Authors:** Saket Choudhary, Rahul Satija

## Abstract

Heterogeneity in single-cell RNA-seq (scRNA-seq) data is driven by multiple sources, including biological variation in cellular state as well as technical variation introduced during experimental processing. Deconvolving these effects is a key challenge for preprocessing workflows. Recent work has demonstrated the importance and utility of count models for scRNA-seq analysis, but there is a lack of consensus on which statistical distributions and parameter settings are appropriate. Here, we analyze 58 scRNA-seq datasets that span a wide range of technologies, systems, and sequencing depths in order to evaluate the performance of different error models. We find that while a Poisson error model appears appropriate for sparse datasets, we observe clear evidence of overdispersion for genes with sufficient sequencing depth in all biological systems, necessitating the use of a negative binomial model. Moreover, we find that the degree of overdispersion varies widely across datasets, systems, and gene abundances, and argues for a data-driven approach for parameter estimation. Based on these analyses, we provide a set of recommendations for modeling variation in scRNA-seq data, particularly when using generalized linear models or likelihood-based approaches for preprocessing and downstream analysis.

## Introduction

Single-cell RNA-sequencing (scRNA-seq) represents a powerful approach for the unsupervised characterization of molecular variation in heterogeneous biological systems (1, 2). However, separating biological heterogeneity across cells that corresponds to differences in cell type and state from alternative sources of variation represents a key analytical challenge in the normalization and preprocessing of single-cell RNA-seq data (3, 4). Upstream analytical workflows typically aim to achieve two separate but related tasks. First, data normalization aims to correct for differences in cellular sequencing depth, which collectively arise from fluctuations in cellular RNA content, efficiency in lysis and reverse transcription, and stochastic sampling during next-generation sequencing (5). Second, variance stabilization aims to address the confounding relationship between gene abundance and gene variance, and to ensure that both lowly and highly expressed genes can contribute to the downstream definition of cellular state. Although the use of unique molecular identifiers (UMIs), random sequences that label individual molecules, has been a promising approach to limit amplification bias (6, 7), variation due to sequencing depth still arises in such datasets and can be a major source of technical variance. These challenges are not unique to single-cell sequencing (8), but the sparsity of scRNA-seq data, coupled with substantial diversity in profiling technologies, necessitates the development and assessment of new methods.

While initial work focused on the development of cell ‘size-factors’ for normalization, recent methods have been focused on the development and application of statistical models for scRNA-seq analysis. In particular, two recent studies proposed to use generalized linear models (GLMs), where cellular sequencing depth was included as a covariate, as part of scRNA-seq preprocessing workflows. Our sctransform (9) approach utilizes the Pearson residuals from negative binomial regression as input to standard dimensional reduction techniques, while GLM-PCA (10) focuses on a generalized version of principal component analysis (PCA) for data with Poisson-distributed errors. More broadly, multiple techniques aim to learn a latent state that captures biologically relevant cellular heterogeneity using either matrix factorization or neural networks (11–13), alongside a defined error model that describes the variation that is not captured by the latent space.

Together, these studies demonstrate the importance and potential of statistical models to assist in the normalization, variance stabilization, and downstream analysis of scRNA-seq data. However, each of these approaches requires an explicit definition of a statistical error model for scRNA-seq, and there is little consensus on how to define or parameterize this model. While multiple groups have utilized a Poisson error model (10, 14–18), others argue that the data exhibit evidence of overdispersion, requiring the use of a negativebinomial (NB) distribution (5, 19–21). Even for methods that assume a NB distribution, different groups propose different methods to parameterize their model. For example, a recent study (22) argued that fixing the NB inverse overdispersion parameter *θ* to a single value is an appropriate estimate of technical overdispersion for all genes in all scRNA-seq datasets, while others (23) propose learning unique parameter values for each gene in each dataset. This lack of consensus is further exemplified by the scvi-tools (11, 24) suite, which supports nine different methods for parameterizing error models. The purpose of this error model is to describe and quantify heterogeneity that is not captured by biologically relevant differences in cell state, and highlights a specific question: How can we model the observed variation in gene expression for an scRNA-seq experiment conducted on a biologically ‘homogeneous’ population?

## Results

### Shallow sequencing masks overdispersion in scRNA-seq data

We first explored whether a Poisson distribution was capable of fully encapsulating heterogeneity in scRNA-seq data that was independent of biological variation in the cellular state (i.e., ‘independent of the latent space’ (25)). The rationale behind a Poisson model assumes that homogeneous cells express mRNA molecules for a given gene at a fixed underlying rate, and the variation in scRNA-seq results specifically from a stochastic sampling of mRNA molecules, for example due to inefficiencies in reverse transcription and PCR, combined with incomplete molecular sampling during DNA sequencing (5, 25). The Poisson distribution constrains the variance of a random variable to be equal to its mean, and has been utilized for modeling UMI counts in multiple previous studies (15, 16). While the Poisson distribution is well suited to capture variation driven by stochastic technical loss and sampling noise, it cannot capture other sources of biological heterogeneity between cells that are not driven by changes in cell state, for example, intrinsic variation caused by stochastic transcriptional bursts (26–28). These fluctuations would cause scRNA-seq data to deviate from Poisson statistics, exhibiting overdispersion that can be modeled using a negative binomial distribution.

We therefore asked whether scRNA-seq data exhibited evidence of overdispersion by exploring the mean-variance relationship using technical controls (endogenous RNA and spike-ins), cell line (HEK293 and NIH3T3), and heterogeneous (PBMC; mouse cortex; fibroblasts) datasets profiled using multiple technologies (Supplementary Table S1). These datasets have varying sequencing depths with median UMIs per cell spanning from approximately 375 to more than 195, 000 (Supplementary Figure S1). In each dataset, we performed a goodness-of-fit test, independently modeling the observed counts for each gene to be Poisson distributed, while accounting for differences in sequencing depth between individual cells (Supplementary Methods). For the technical control datasets (8, 14), where the input to each ‘cell’ represented a uniform source of RNA, observed variation was largely consistent with the Poisson model (Figure 1B). In contrast, when analyzing a human PBMC dataset profiled using Smart-seq3 (29), thousands of genes were poorly fit by a Poisson distribution (Figure 1A and B), even after accounting for cell-to-cell variation in sequencing depth (Supplementary Table S2). While we expected to observe overdispersion for a subset of genes, particularly for those whose expression varies across multiple cell types, we were surprised to see that 97.6% of genes with average expression *>* 1 UMI/cell failed the Poisson goodness-of-fit test. We observed a similar phenomenon when analyzing data from homogeneous HEK293 cells profiled with the 10X Chromium v2 system (HEK-r2; Figure 1A and B), with 93% of genes exhibiting average abundance of *>* 1 UMI/cell demonstrating evidence of overdispersion. In each of the 58 datasets we analyzed, genes exhibiting Poisson variation were overwhelmingly lowly expressed compared to genes that were overdispersed (Supplementary Figure S2). Moreover, when comparing results for cell-line datasets where we expect low levels of variation in cell state, we found that the global fraction of genes deviating from a Poisson distribution was correlated with the average sequencing depth of the dataset (Figure 1C).

**Figure 1.**
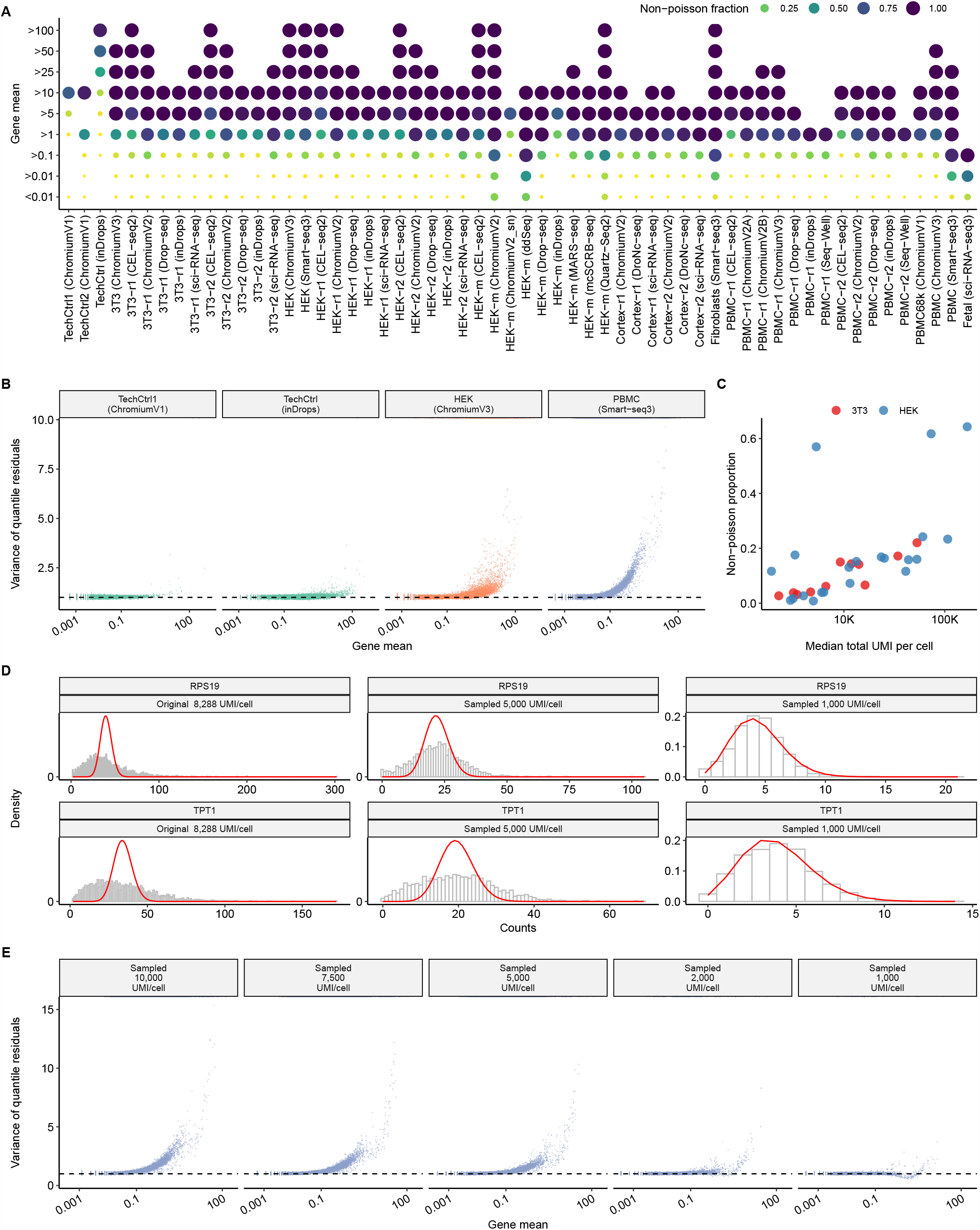
Shallow sequencing masks overdispersion in scRNA-seq data. **A)** Proportion of genes that fail a goodness-of-fit test for a Poisson GLM (Supplementary Methods), as a function of gene abundance, for 58 scRNA-seq datasets. For visual clarity, both the color and diameter of each dot correspond to the fraction of genes that exhibit overdispersion. Y-axis represents non-cumulative gene abundance bins between two consecutive labels (for example, *>* 1 refers to all genes with average abundance *>* 1 UMI and *≤* 5 UMI). Values are listed in Supplementary Table S2. **B)** Relationship between average gene abundance and quantile residual variance, after applying a Poisson GLM (Supplementary Methods). Results are shown for datasets profiling endogenous RNA (‘technical controls’), a HEK293 cell line (‘biological controls’), and human PBMC (‘heterogeneous’). **C)** In datasets profiling cell lines, the fraction of genes that exhibit overdispersion is correlated with average sequencing depth. **D)** Distribution of molecular counts for highly expressed genes in the PBMC Smart-seq3 dataset after downsampling to two different sequencing depths. The expected density assuming a Poisson distribution is shown in red. **E)** Same as (A) but after downsampling the PBMC Smart-seq3 dataset to five different sequencing depths.

Our results suggest that scRNA-seq datasets commonly exhibit biological variation that exceeds Poisson sampling, but that the statistical power to detect these fluctuations requires sufficient sequencing depth. For example, when observing molecular counts in the deeply sequenced PBMC dataset (median 8, 288 UMI/cell), highly expressed genes such as TPT1, RPS19 exhibited particularly strong deviations from Poisson variability (Figure 1D). However, we found that when artificially downsampling the same dataset to 1, 000 UMI/cell, a depth that is common to shallowly sequenced scRNA-seq datasets, deviations from a Poisson distribution were strongly reduced (Figure 1E). After downsampling, only 0.5% genes failed the Poisson goodness-of-fit test, demonstrating that reducing cellular sequencing depth can artificially create the appearance of Poisson variation. We conclude that the Poisson error model may represent an acceptable approximation for scRNA-seq datasets with shallow sequencing, but as the sensitivity of molecular profiling continues to increase, error models that allow for overdispersion are required for scRNA-seq analysis. Furthermore, we reiterate that the use of a Poisson error model does not account for the possibility of intrinsic stochastic noise in single-cell datasets, though this type of noise has been extensively described and does not correlate with changes in cell type or state.

### The level of overdispersion varies substantially across datasets

We next focused on the application of negative binomial error models, and considered different strategies for parameterizing the level of overdispersion associated with each gene. Recent work (22) suggested that a negative binomial model with a fixed parameterization (for example, inverse overdispersion parameter *θ* = 100) could be applied to all scRNA-seq datasets to achieve effective variance stabilization. To explore whether a single value of *θ* could be applied to diverse scRNA-seq datasets, we first independently fit *θ* estimates for each gene in each dataset using a GLM with negative binomial errors (NB GLM), using library size as an offset to account for variation in cellular sequencing depth. We observed substantial differences in the magnitude of the estimated *θ* across different datasets, though replicate datasets from the same study yielded concordant results (Figure 2A and B). Consistent with our previous results (Figure 1B), *θ* values for each dataset varied across different biological systems, technologies, and sequencing depths.

**Figure 2.**
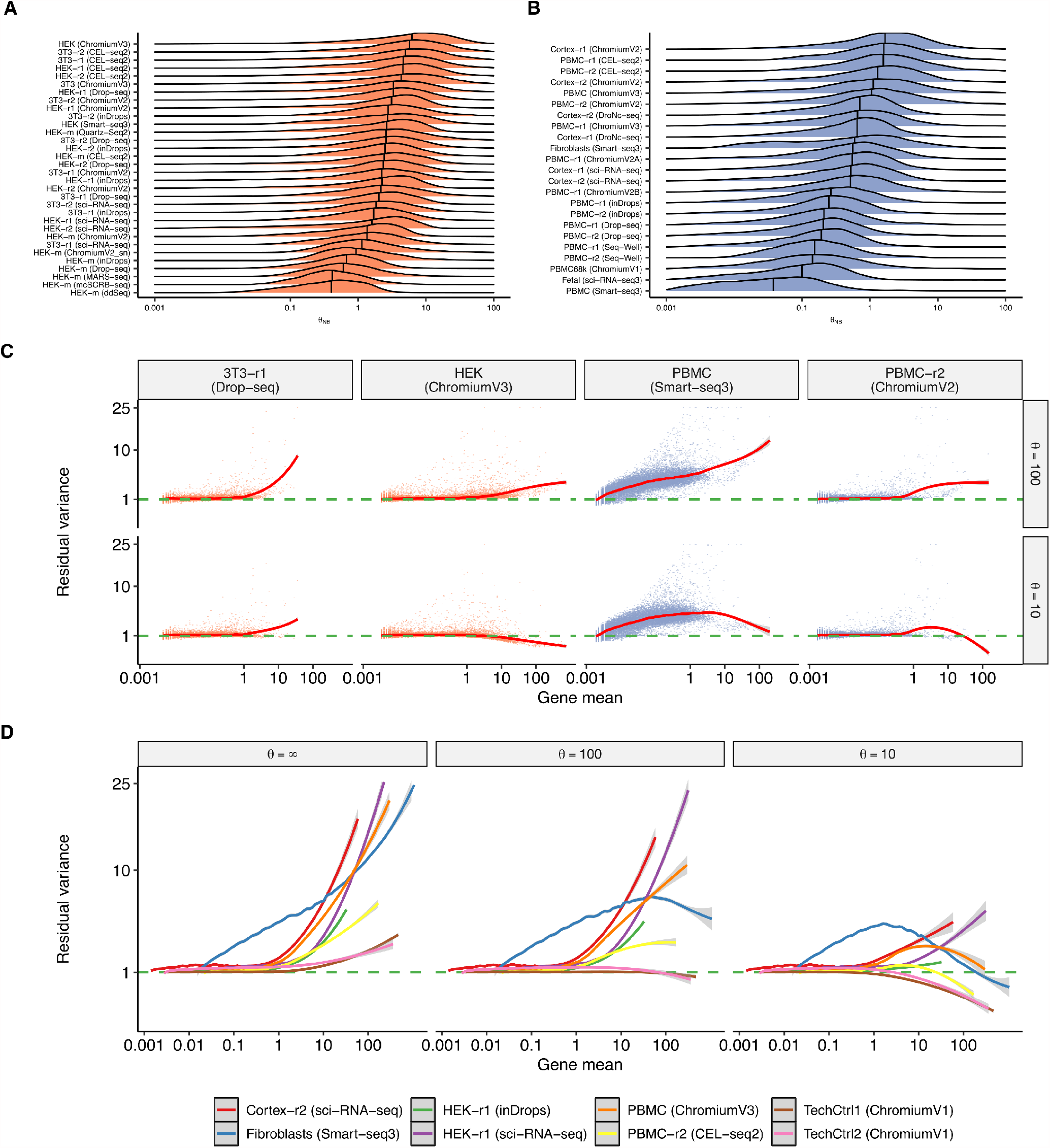
Overdispersion varies across datasets. **A), B)** Distribution of per-gene values for the estimated inverse overdispersion *θ*_*NB*_ of a NB GLM across a range of cell lines (A) and heterogeneous datasets (B). We estimated parameters only for genes where the variance of counts exceeds the mean. Vertical bar indicates the median of the distribution, which varies substantially across datasets, but is concordant across replicate experiments within the same study. **C)** Relationship between gene mean and the variance of Pearson residuals resulting from an NB GLM with *θ* = 10 or *θ* = 100. Each dot is a gene and the trendline (LOESS) is shown in red. **D)** Same as (C), but shown for additional datasets and for *θ* = *∞* (Poisson). Only trendlines are shown for visual clarity.

We next tested the ability for a single value of *θ* to perform effective variance stabilization across a range of datasets. We processed each of our 58 datasets using an NB GLM after fixing *θ* to a single value for all genes in the dataset (for example, *θ*=100). We found that no single value of *θ* could achieve effective variance stabilization across all datasets. For example, a negative binomial error model with *θ* = 100 resulted in clear heteroskedasticity in multiple datasets (Figure 2C), as we observed a strong relationship between the mean expression of a gene, and its residual variance. This will artificially boost the weight of all highly expressed genes in downstream analysis such as dimensional reduction and clustering. We repeated the analysis with two alternative models, setting *θ* = ∞ and *θ* = 10, both of which revealed similar shortcomings in multiple datasets (Figure 2D and Supplementary Figures S3 - S10). We conclude that fixing a single value of *θ* may achieve effective performance in certain cases, but is unlikely to generalize across the diversity of systems and technologies represented by scRNA-seq data.

### Gene overdispersion varies as a function of abundance

An alternative strategy for parameterizing *θ* leverages a well-characterized strategy for modeling counts in bulk RNA-seq data, where per-gene dispersion estimates have repeatedly been found to vary as a function of expression abundance (30–36). In sctransform (9), we aim to estimate a global relationship between gene abundance and *θ* by employing a regularization procedure where parameters are first fit for each gene individually, but information from genes with similar average abundances is subsequently pooled together in order to improve the robustness of parameter estimates. The underlying rationale for this choice is the non-decreasing relationship between gene abundance and *θ* that has been repeatedly observed in bulk RNA-seq studies (30–36). When analyzing each of the technologies and biological systems explored in this manuscript, we identified the same global patterns relating gene abundance and overdispersion levels (Supplementary Figures S11 - S14).

We also considered the findings from (22), which proposed that *θ* values should not vary as a function of gene abundance, and suggested that the relationship between these two variables was driven entirely by biases in the parameter estimation procedure, especially when analyzing lowly expressed genes. We first confirmed that lowly expressed genes, particularly those with average abundance *<* 0.1 UMI/cell, posed difficulties for parameter estimation. This is because the vast majority of count values for these genes are 0, creating inherent challenges in maximum likelihood estimation. When estimating parameters on simulated data drawn from a negative binomial with fixed *θ*, we confirmed a bias for these genes that resulted in decreased parameter estimates for *θ* (Supplementary Figure S15). However, using two complementary analyses, we found that this bias was not sufficient to explain the true relationships we observed in biological data. First, we observed a non-decreasing relationship between gene abundance (*µ*) and dispersion (*θ*) even when moving beyond the threshold for lowly expressed genes, which we did not observe when analyzing simulated data (Supplementary Figure S16). Additionally, we attempted to increase (‘up-sample’) the depth of single cell datasets by pooling together molecular counts from cells with similar molecular profiles (Supplementary Methods) as inspired by the MetaCell framework (37). We repeated the parameter estimation procedure on metacells generated either from single-cell data, or using our simulation framework (Methods). Increasing the depth of sampling removed the effects of bias when analyzing simulated data, but preserved the observed relationship between *µ* and *θ* on real biological datasets (Supplementary Figure S16). We conclude that when modeling scRNA-seq data using a negative binomial distribution, the inverse overdispersion parameter *θ* does vary as a function of gene abundance, but the true nature of this relationship can be masked for genes with low molecular counts.

### Recommendations for modeling heterogeneity in scRNA-seq datasets

Our findings highlight how the extensive diversity of scRNA-seq datasets poses challenges in identifying uniform procedures for the preprocessing and normalization of scRNA-seq data. Sparsely sequenced datasets may appear to be compatible with the use of Poisson error models, but datasets with additional sequencing depth reveal clear evidence of overdispersion. The level of overdispersion, exemplified by the NB parameter *θ*, also can vary substantially across datasets, technologies and systems, and even varies within a dataset as a function of gene abundance. However, the estimation of robust parameter estimates for *θ* can be challenging for lowly expressed genes, especially when analyzing datasets with sparse sequencing. We therefore considered recommendations for balancing these considerations, providing the flexibility to learn error models that can be robustly applied to our full spectrum of scRNA-seq datasets.

We first recommend the use of negative binomial observation model as an alternative to the Poisson distribution. Our analyses show that the Poisson distribution is a reasonable approximation for technical-control datasets consisting of endogenous or spike-in RNA, as well as for some scRNA-seq experiments with shallow sequencing. However, scRNA-seq datasets from cell lines exhibit clear evidence of overdispersion at higher sequencing depths, even for genes that do not correlate with changes in cell type or state. At least some of this overdispersion likely originates from ‘intrinsic’ noise, stochastic cellular variation that is inherent to the processes of mRNA transcription and degradation, and will affect the expression heterogeneity of all genes. While this variation is not a result of measurement error, it is not the primary focus of downstream scRNA-seq analyses, including the identification of cell types and states, and the inference of developmental trajectories. We therefore recommend that this variation be modeled independently of the latent space, which requires the use of a negative binomial error model. We note that the Poisson distribution is a special case of the negative binomial, and therefore the NB model can be successfully applied for datasets with very shallow sequencing, with appropriate parameter settings.

Second, we recommend learning negative binomial parameters separately for each dataset, rather than fixing them to a single value across all analyses. Moreover, we recommend allowing *θ* to vary across genes within a dataset, as a function of average gene abundance. The relationship between *µ* and *θ* has been repeatedly demonstrated in bulk RNA-seq, and is apparent across diverse scRNA-seq datasets as well, particularly for genes with sufficient sequencing depth. Using a fixed *θ* to parameterize all genes in a scRNA-seq dataset leads to ineffective variance stabilization and results in a global relationship between expression level and expression variance (Figure 2 and Supplementary Figures S3 and S4). We note that the recommendations described above relate not only to GLM-based preprocessing workflows, but also probabilistic or likelihood-based models (11, 24, 38).

Our analyses highlighted that lowly expressed genes with particularly sparse molecular counts often lacked sufficient information content to robustly detect overdispersion. We therefore designed a modified regularization procedure for learning GLM parameter estimates (Supplementary Methods). First, following the recommendations from (22), we fix the slope of the NB GLM to its analytically derived solution of ln(10), so that only the overdispersion and intercept parameters are estimated for each gene. Second, we reasoned that for genes with very low expression (*µ <* 0.001), or where the variance of their molecular counts does not exceed the mean (i.e. *σ*^2^ ≤ *µ*), we do not have sufficient evidence for overdispersion to fit negative binomial parameters. We therefore removed these genes from the regularization process and fixed their *θ* parameter to ∞, exemplifying a Poisson distribution. For example, in the scRNA-seq dataset of HEK cells profiled with SMART-Seq3, we removed 1, 577 genes (8.5%) at this stage, the majority of which were lowly expressed (66.64% *<* 0.1 UMI/cell). We found that our modified regularization enables us to reproducibly learn gene-specific parameters even when using a subset of cells in the estimation procedure. This indicates increased robustness (Figure 3A), and allows us to learn a regularized relationship between *µ* and *θ* using only a subset of cells that achieves nearly identical results (Figure 3B) with increased computational efficiency (Figure 3C).

**Figure 3.**
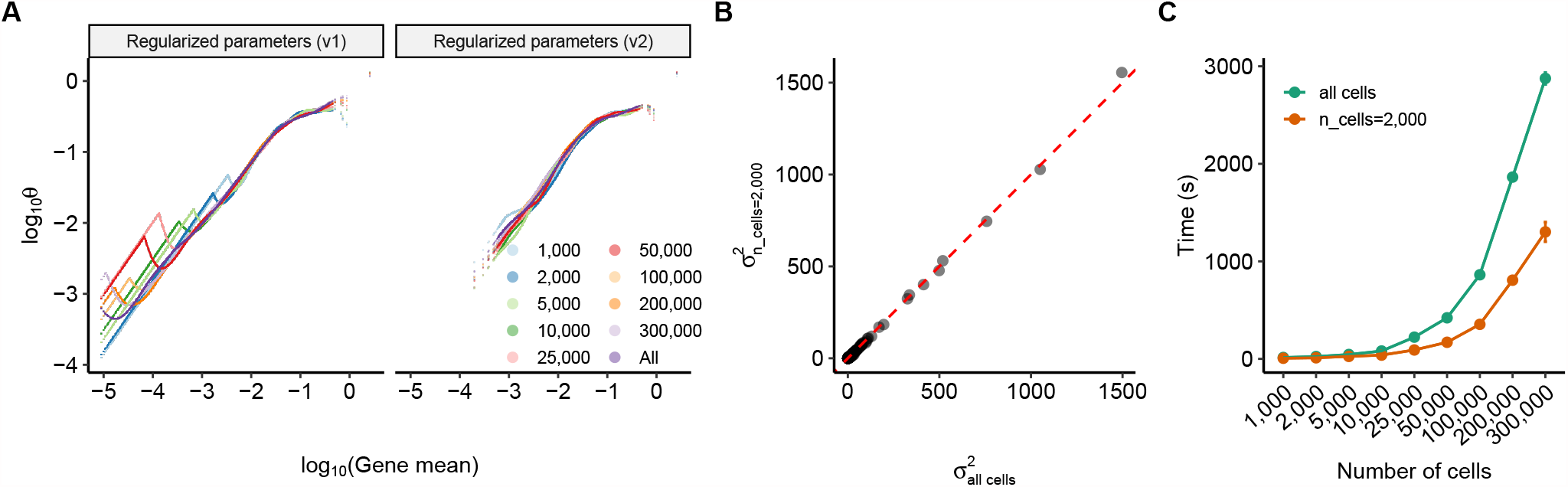
A modified regularization procedure improves the robustness of parameter estimates. **A)** Left: Estimated parameter estimates for *θ* on the Fetal sci-RNA-seq3 dataset (39), using the original regularization procedure from (9) (v1 regularization). Regularized estimates were learned using all cells (purple line), or downsampled cell subsets. Right: Same as (A), but using a modified procedure where the GLM slope was fixed, and genes where *σ*^2^ *≤ µ* and *µ <* 0.001 were excluded from regularization (v2 regularization) which improves robustness, and enables us to learn parameter estimates from a subsample of 2, 000 cells. **B)** Correlation of Pearson residual variance after applying a NB GLM with v2 regularization where parameters were estimated from all 377, 456 cells (x-axis), and a subsample of 2, 000 cells (y-axis). **C)** Green curve: total sctransform run time as a function of dataset size, using all cells to estimate parameters. Orange curve: total runtime when using a subsample of 2, 000 cells, which increases computational efficiency for large datasets.

To test the broad applicability of this procedure, we applied it to each of the 58 datasets examined in this manuscript. In each case, we achieved effective variance stabilization as we observed no global relationship between gene expression levels and the variance of the resulting Pearson residuals (Supplementary Figures S17 - S20). Moreover, in each case, genes with the highest residual variance were distributed across a range of expression levels and - when analyzing heterogeneous samples - represented markers that have been strongly associated with individual cell types. As a result, application of this preprocessing pipeline will give the greatest weight to these markers, while downweighting fluctuations in the most highly expressed genes, which often appear to exhibit extensive heterogeneity in the absence of variance stabilization. These results indicate that our preprocessing workflow has sufficient flexibility to accurately model a wide variety of scRNA-seq datasets and serves as a basis for our recommendations in this manuscript.

## Discussion

The application of statistical count models for preprocessing of scRNA-seq data overcomes important challenges that cannot be addressed by using linear size or scaling factor-based normalization. However, these techniques require the selection of an appropriate error distribution and accompanying parameter settings. Here, we explore these questions through the analysis of a wide diversity of scRNA-seq datasets varying across technologies, biological systems, and sequencing depths.

Our analyses revealed three key insights. First, we found that all scRNA-seq datasets exhibited clear evidence of overdispersion (i.e. deviation from a Poisson distribution), even after accounting for differences in sequencing depth, once exceeding a minimum expression level. This threshold varied across datasets as a function of average sequencing depth. This result strongly supports the use of negative binomial error models when analyzing UMI datasets. Second, we found that the negative binomial overdispersion parameter *θ* varied substantially across datasets, arguing against the use of a fixed *θ* estimate. Finally, we found that all datasets exhibited a dependence between gene abundance and overdispersion estimates. This result is robust even when considering potential biases in the overdispersion parameter estimation, and supports an empirical approach to learn regularized parameter estimates, as is commonly performed in bulk RNA-seq analysis.

Taken together, these results are compatible with the idea that cell-to-cell variation in scRNA-seq count data can be decomposed into multiple broad categories. The first represents variation in cell type and state which is biologically driven and encoded in cellular transcriptomes. This heterogeneity is the primary interest and focus of downstream analysis, and is typically represented in a latent space that can be learned via linear or non-linear dimensional reduction techniques. A second source represents technical measurement error arising from the stochastic loss of molecules during library preparation and sequencing. This sampling error can be modeled using a Poisson distribution and, particularly for shallowly sequenced datasets, represents a substantial source of remaining heterogeneity.

Our analyses suggest a third level of variation that should be accounted for: fluctuations in gene expression which are driven by the noise that is inherent to the processes of mRNA transcription and degradation (i.e. ‘intrinsic noise’) and manifests as overdispersion in scRNA-seq data. The presence of intrinsic noise has been extensively characterized and is an inevitable consequence of the gene regulatory process. Therefore, no two cells can generate mRNA molecules at exactly the same rate (an assumption of a Poisson process), even if they originate from the same ‘homogeneous’ population. Our analyses demonstrate that intrinsic noise is readily detectable for genes with sufficient sequencing depth, but can be masked in shallow datasets (Figure 1E). While intrinsic noise it is not driven by measurement error, it should also be modeled independently of the latent space. Therefore, as the sensitivity and depth of scRNA-seq datasets continue to increase, the use of negative binomial error models will become increasingly important. Moreover, the level of intrinsic noise can vary across biological systems and gene abundance levels, motivating the use of a data-driven regularization procedure to learn gene-level overdispersion parameters.

Our analyses highlight the importance of considering a diversity of datasets when evaluating the statistical properties of new data types. While our results are therefore applicable to scRNA-seq measurements, they cannot be directly applied to new single-cell modalities, including protein measurements (i.e. CITE-seq (40)), chromatin accessibility profiles (i.e. scATAC-seq (41)), and DNA interaction maps (i.e. scCUT&TAG (42, 43)). As with cellular transcriptomes, these modalities can be profiled using multiple technologies that vary in their sensitivity and sparsity. We anticipate exciting future work that will characterize and parameterize heterogeneity in these data types, to achieve effective preprocessing and normalization.

### Availability of materials and data

Raw datasets used in the main text are available from public URLs listed in Supplementary Table S1. Scripts to reproduce the analyses are available at https://github.com/saketkc/scRNA_NB_comparison.

Source code for sctransform along with the modifications described in this manuscript is available in the forked repository at https://github.com/saketkc/sctransform. A Python implementation that interfaces with the Scanpy (44) package is available at https://github.com/saketkc/pysctransform.

To invoke ‘v2’ regularization in `SCTransform` using Seurat (45):

~~~
library(Seurat)
object <- CreateSeuratObject(counts = counts)
object <- SCTransform(object, vst.flavor=‘v2’)
~~~

Analogously, to use SCTransform in Python (using Scanpy (44)):

~~~
from pysctransform import SCTransform
adata = sc.read_h5ad(“anndata_object.h5ad”)
residuals = SCTransform(adata, vst_flavor=‘v2’)
~~~

## Acknowledgements

The authors would like to thank Christoph Hafemeister, and members of the Satija Lab for thoughtful discussions related to this work. This work was supported by the Chan Zuckerberg Initiative (EOSS-0000000082, HCA-A-1704-01895), and the NIH (RM1HG011014-02, 1OT2OD026673-01, DP2HG009623-01) to R.S.

## Supplementary Methods

### Data sources and preprocessing

All datasets were obtained as preprocessed count matrices from Gene expression omnibus (GEO), EBI ArrayExpress, or author’s website. In each case, we utilized cells that had passed the QC thresholds set by the original study authors. However, to minimize the effect of potential cell outliers in our data, we filtered out cells that fell outside of the 5% to 95% UMI quantiles in each dataset. Additionally, we removed all cells where more than 15% of reads mapped to mitochondrial transcripts. We did not perform any filtering for the Fetal sci-RNA-seq3 dataset as it had already been filtered and annotated by the authors. The dataset source and associated publication are available in Supplementary Table S1.

### Goodness of Fit test using a Poisson GLM

To explore whether a Poisson distribution represents an appropriate error model for UMI-based scRNA-seq data, we fit a Poisson GLM adjusting for differences in library size modeled as an offset. To reduce the computational complexity, we subsampled 1, 000 cells in a density dependent manner from the count matrices of each dataset: the probabilty of selecting a cell *c* is 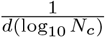, where *d* is the density estimate of all log_10_-transformed total cell UMIs and *N*_*c*_ is the total UMI counts in cell *c*. These subsampled count matrices were then used to fit a Poisson GLM for each gene UMI vector with total UMI content of each cell modeled as an offset vector (`glm.fit(gene_umi ∼ 1, offset=log(total_umi), family=Poisson(link=“log”)`) in `R`. We then performed a goodness of fit test on the randomized quantile residuals (46) of this GLM model fit calculated using `statmod::qresid(model)`. If the data is well-described by the model, the sum of squares of the quantile residuals is expected to follow a Chi-squared distribution with degrees of freedom = *N*_cells_ − 1 where *N*_cells_ represents the total number of cells in the dataset. We chose quantile residuals to measure the goodness of fit, as they have lower type-I error and higher power in comparison to other residuals for identifying misspecification (47). To calculate p-values, we used the `pchisq` function in R (`pchisq(qresid, df=model$df.residual, lower.tail=FALSE)`). To control for multiple testing, we adjusted p-values using the `qvalue` method available through the qvalue package (48). We used a q-value threshold of 0.01 to accept or reject the fit to the Poisson model. Library sizes reflected in Figure 1E were calculated based on the subset count matrices.

### Assessing overdispersion after downsampling sequence depth

In Figure 1D-E we assess the level of dispersion in the PBMC Smart-seq3 dataset, after downsampling the dataset to different sequencing depths. The full dataset contains 2, 629 cells with a median UMI/cell of 8, 288 with a maximum coverage of 20, 463 UMI/cell. When downsampling to 10, 000 UMI/cell, we first excluded 1, 837 cells where *<* 9, 900 UMIs were detected in the dataset. For the remaining cells, we randomly sampled molecules at a proportion expected to yield 10, 000 UMI/cell on average and retained only cells that contained UMIs in the range 10, 000 ± 100 to minimize the effect of differences in sequencing depth. We repeated this process for multiple sequencing depths shown in Figure 1D-E.

### Comparing levels of overdispersion across datasets

In Figure 2A-B, we fit NB GLM to each gene in each dataset, in order to estimate the inverse overdispersion parameter *θ*. We model the observed counts for each gene using the following model `gene_umi ∼ 1`, and estimate parameters using `glmGamPoi::glm_gp(gene_umi, model, offset=log(total_umi), size_factors=FALSE)` using the glmGamPoi package (49). We perform this procedure for all genes where the variance of the observed counts exceeds the mean.

### Modeling scRNA-seq datasets with sctransform

For clarity, we restate the modeling framework used in sctransform (9). In sctransform, UMI counts across cells in a dataset are modeled using a generalized linear model (GLM). The total UMI count per cell is used as an offset in the GLM. Thus, for a given gene *g* in cell *c*, we have

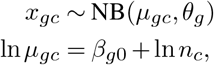

where *θ*_*g*_ is the gene-specific dispersion parameter, *n*_*c*_ = *g x*_*gc*_ is the total sequencing depth and the variance of the negative binomial (NB) is given by 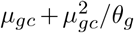.

We perform three steps to remove technical noise and perform variance stabilization. In the first step, the inverse overdispersion parameter *θ* is individually estimated using a subset of genes (2000 by default), which are sampled in a density-dependent manner according to their average abundance. In the next step, we calculate a smoothed curve that characterizes the global relationship between *µ* and *θ*, thereby regularizing *θ* estimates as a function of gene mean. We perform the same regularization for the intercept parameter. We use the geometric mean to estimate gene abundance, which is more robust to outlier values in scRNA-seq. As outlier values can originate from multiple sources including the presence of cell doublets, errors in UMI collapsing, or ambient RNA, we have empirically improved performance when using the geometric mean instead of the arithmetic mean. Although sctransform supports multiple estimators for *θ*, we recommend the use of glmGamPoi (49), an alternate estimator that is more robust and faster.

In the final step, we use the regularized parameter estimates to calculate Pearson residuals *Z*_*gc*_. For each gene-cell combination, the Pearson residuals *Z*_*gc*_ are given by

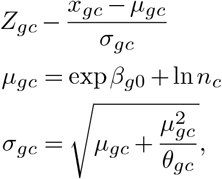

The ‘residual variance’ for a gene represents the remaining variation in gene expression that is not explained by the sctransform model, and is defined as:

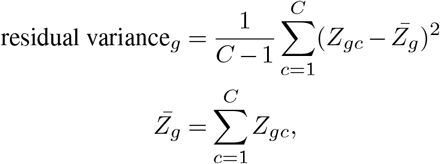

where *C* represents the number of total cells in the dataset.

### Evaluating the performance of a GLM with fixed *θ*

In Figure 2C-D, as well as associated supplementary figures (S3 - S10) we model each of the scRNA-seq datasets using a NB GLM with a fixed value of *θ* for each gene in each dataset. To test this, we utilize the ‘offset’ model as described by Lause et al. in (22). We repeated the analysis with three different values for the fixed overdispersion parameter, *θ* = ∞, *θ* = 100, and *θ* = 10.

### Improving the robustness of parameter regularization

In Figure 3 we explore a modified regularization procedure to improve the robustness of NB parameter estimates, particularly for lowly expressed genes, and to increase computational efficiency. We make two changes to the estimation procedure described in (9). First, we fix the slope parameter of the GLM to ln(10) with log_10_(total UMI) used as the predictor. As described in (22), this value represents the analytically derived solution for this parameter, and closely mirrors the regularized values we had obtained for the slope parameter in the original sctransform procedure. While (22) also recommends fixing the intercept parameter for the GLM, an approximate solution to the maximum likelihood estimate of this parameter can only be obtained for large values of *θ*. As our data-driven estimates for *θ* demonstrate that this parameter can vary substantially across datasets, we continue to set the intercept parameter for the GLM through regularization.

As a second modification, we remove a subset of genes prior to performing regularization. In particular, we reasoned that for genes with either very low abundance (*µ <* 0.001), or where the variance of count values did not exceed the average abundance (i.e. *σ*^2^ ≤*µ*), we lacked sufficient information to learn robust NB parameter estimates. We therefore exclude these genes from the regularization procedure, and set their *θ* parameter estimates to ∞ for all downstream analyses. We note that this filtration step occurs rapidly, as the per-gene mean and variance can be efficiently calculated. For this filtration step, we use the arithmetic mean to set abundance, as this value should be compared with gene variance to determine evidence of overdispersion. For these genes, the regularized intercept 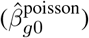 is set to the analytically derived solution from (22), with a fixed slope of ln(10):

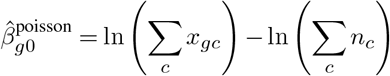

### Simulation of UMI counts with fixed overdispersion

To explore the potential bias of maximum-likelihood (ML) estimators, we simulated an scRNA-seq dataset with fixed levels of overdispersion. We fixed *θ* to different values {0.001, 0.01, 0.1, 1, 10, 100}, and simulated scRNA-seq counts from an NB distribution, using gene means that were taken from the PBMC Smart-Seq3 dataset. We next estimated parameter values for *θ* using both the v1 and v2 versions of our sctransform regularization procedure using glmGamPoi (49) as an estimation engine. We also estimated a maximum likelihood of *θ* using glmGamPoi without explicitly accounting for library size (MLE). To create an ‘upsampled’ dataset where the sequencing depth is higher, we multiplied the estimated means *x*_*gc*_ by a factor of 500, and repeated the sampling procedure.

### Increasing sequencing depth by creating metacells

In order to ‘upsample’ the PBMC Smart-seq3 dataset, we ran MetaCells v0.3.5 (37) for three different values of ‘K’ parameter (200, and 300, and 400) with all other parameters as defaults. UMI counts of cells belonging to one metacell were consolidated to create a metacell count, resulting in a higher sequencing depth. These metacells were then used as input to sctransform to estimate per gene *θ*.

## Supplementary Figures & Tables

**Figure S1.**
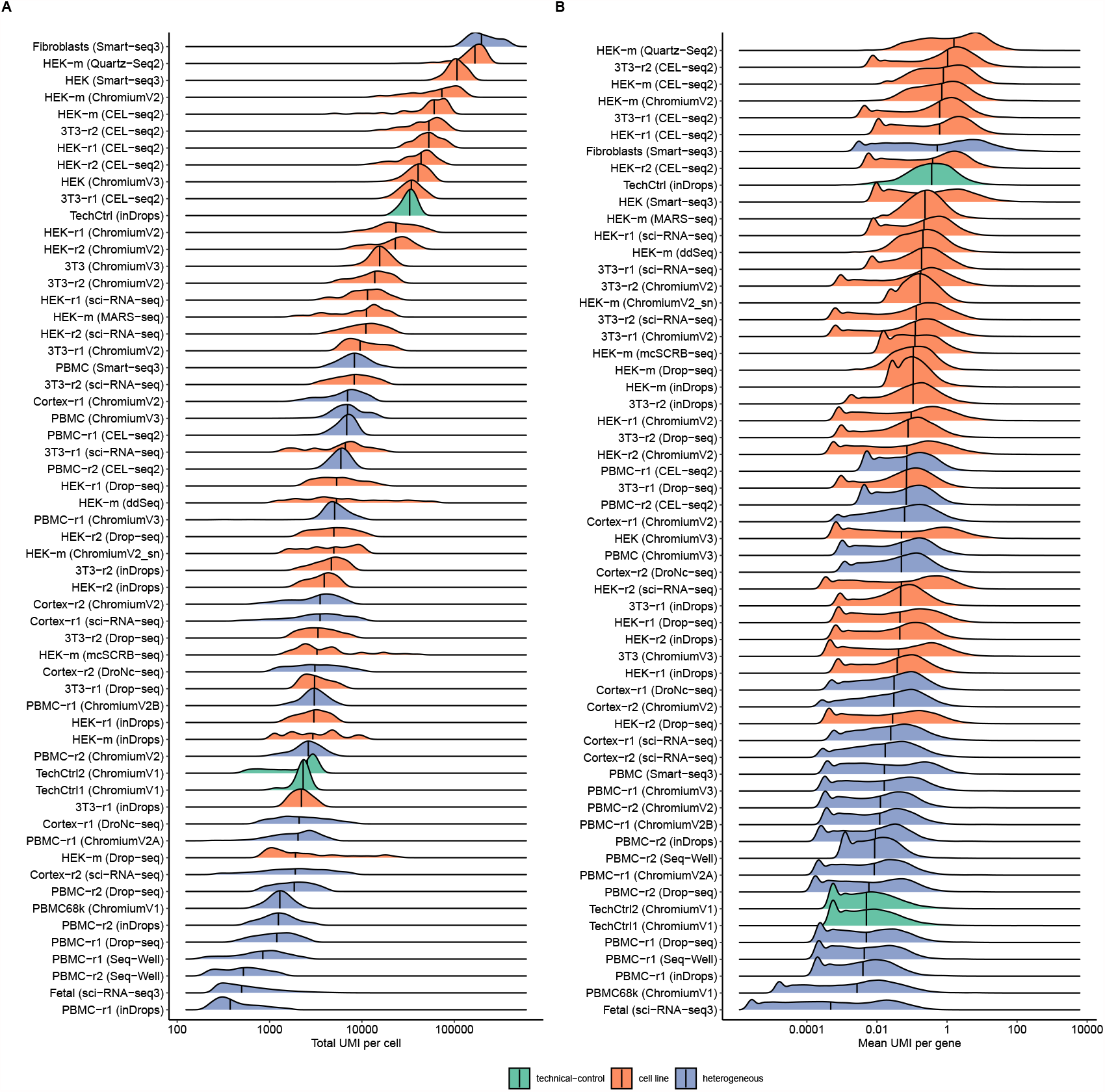
UMI statistics for the 58 datasets analyzed in this manuscript. **A)** Distribution of total UMI per cell across datasets **B)** Distribution of mean UMI per gene across datasets (technical control = endogeneous or spike-in RNA; cell line = HEK293 and 3T3 cell lines; heterogeneous = samples consisting of multiple cell types).

**Figure S2.**
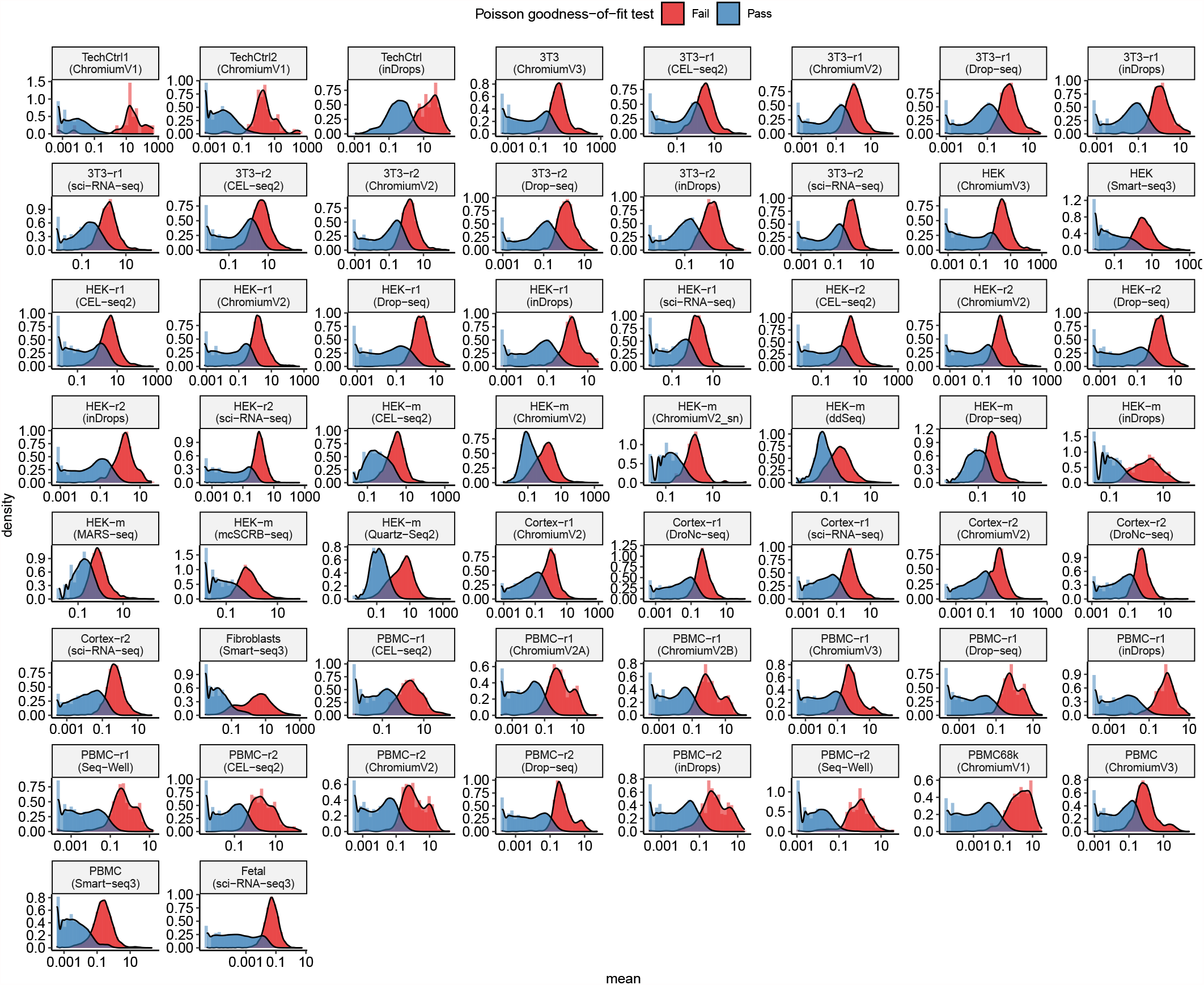
Genes that exhibit Poisson heterogeneity are more lowly expressed. In each dataset, we performed a per-gene goodness-of-fit test based on a GLM with a Poisson error model (Supplementary Methods). Shown are the distribution of gene abundances (average UMI/gene) for genes that passed (blue) and failed (red) the goodness-of-fit test.

**Figure S3.**
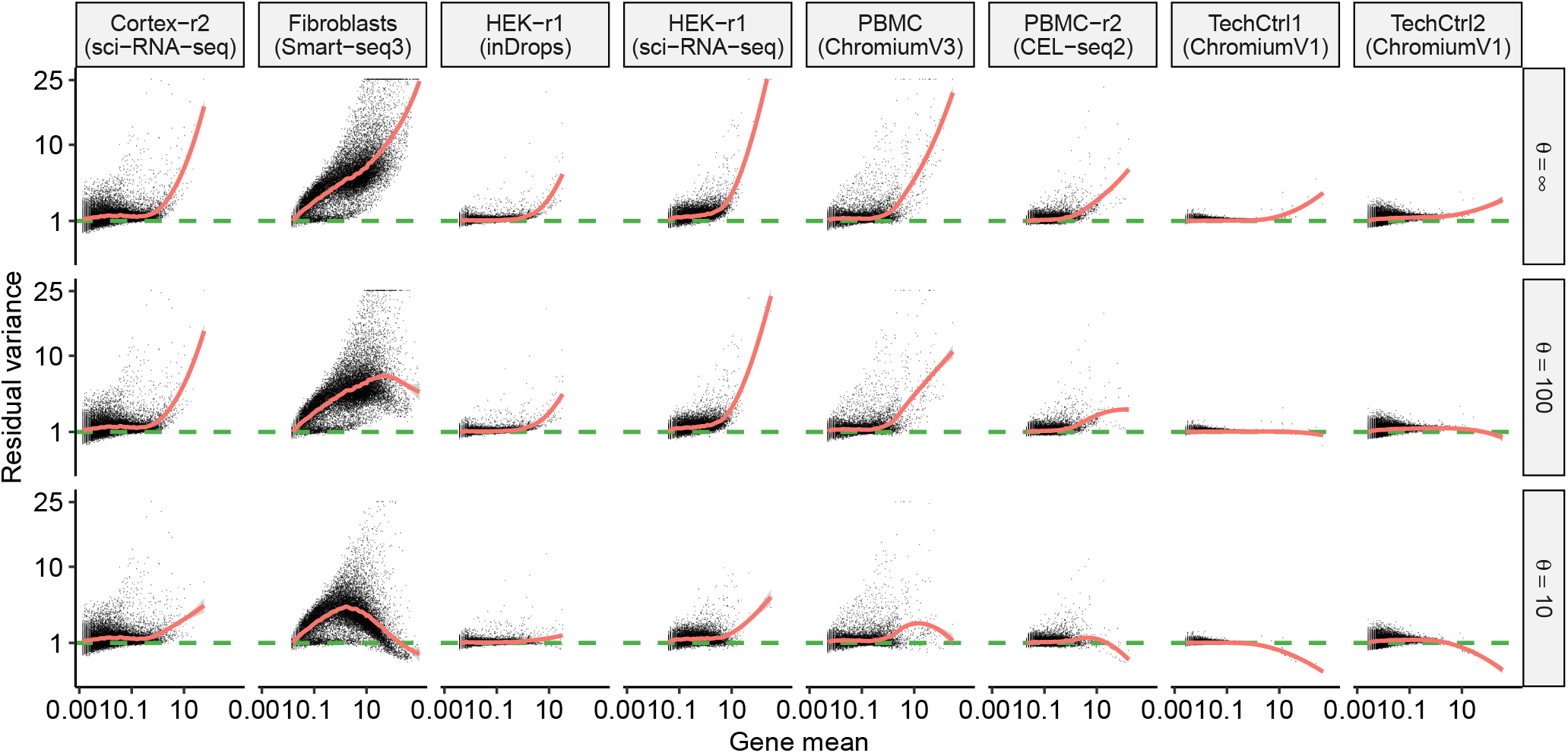
Relationship between gene abundance and the variance of Pearson residuals. Values shown are resulting from an NB GLM with three different values of *θ*. Same as Figure 2D but with per-gene estimates highlighted instead of smoothed curves.

**Figure S4.**
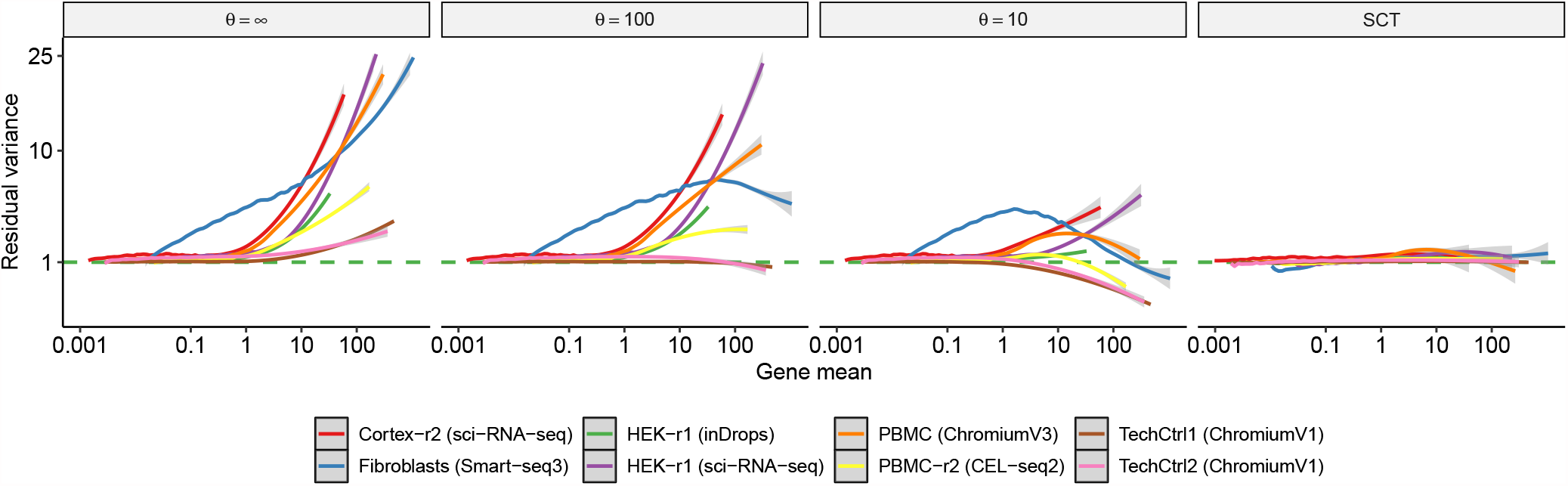
Relationship between gene abundance and the variance of Pearson residuals. Same as Figure 2D but additionally showing results for sctransform (v2 regularization).

**Figure S5.**
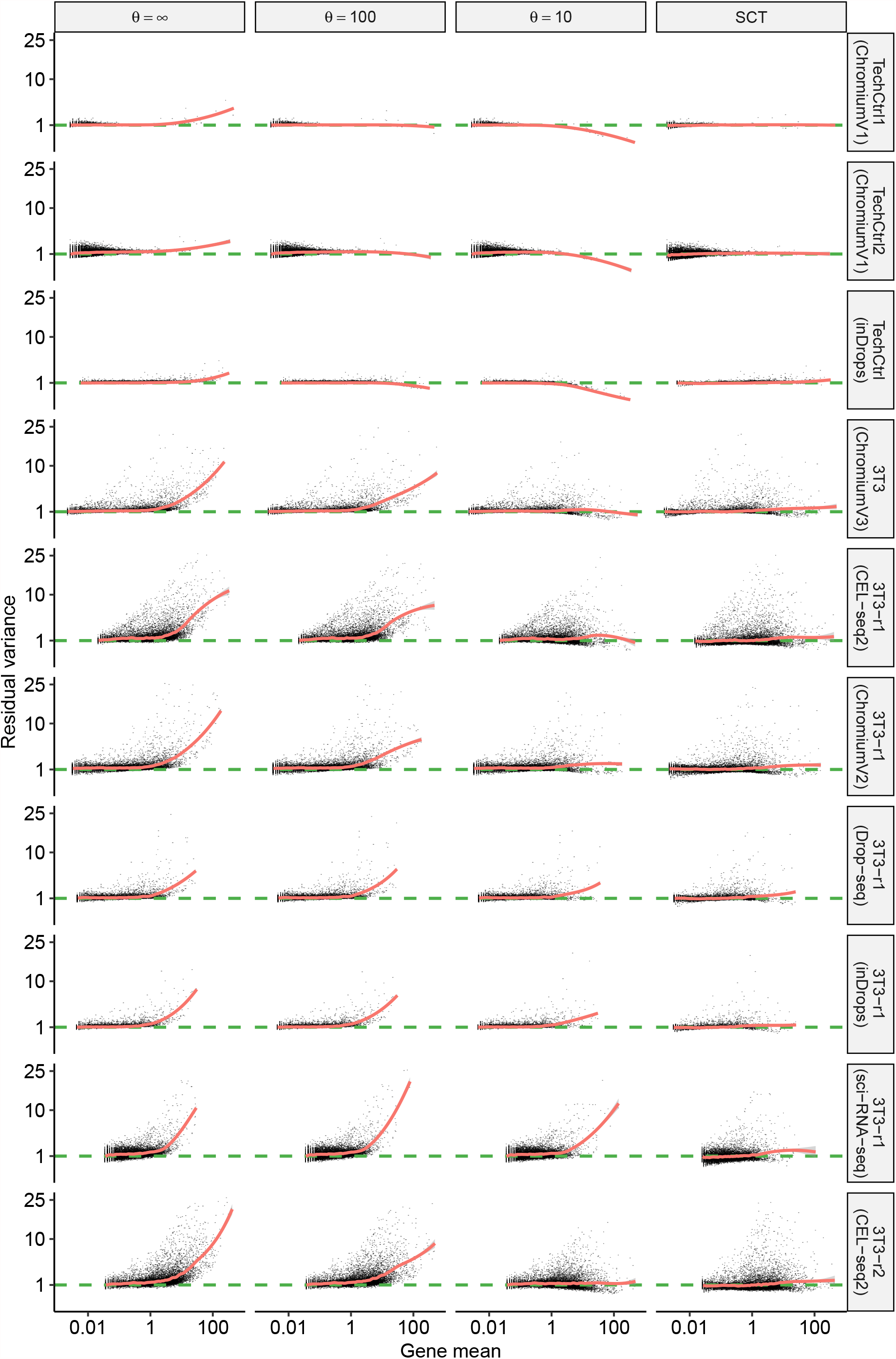
Evaluating variance stabilization for different error models. Same as in Figure 2C, but with additional datasets 1-10. Also shown are results from sctransform (v2 regularization).

**Figure S6.**
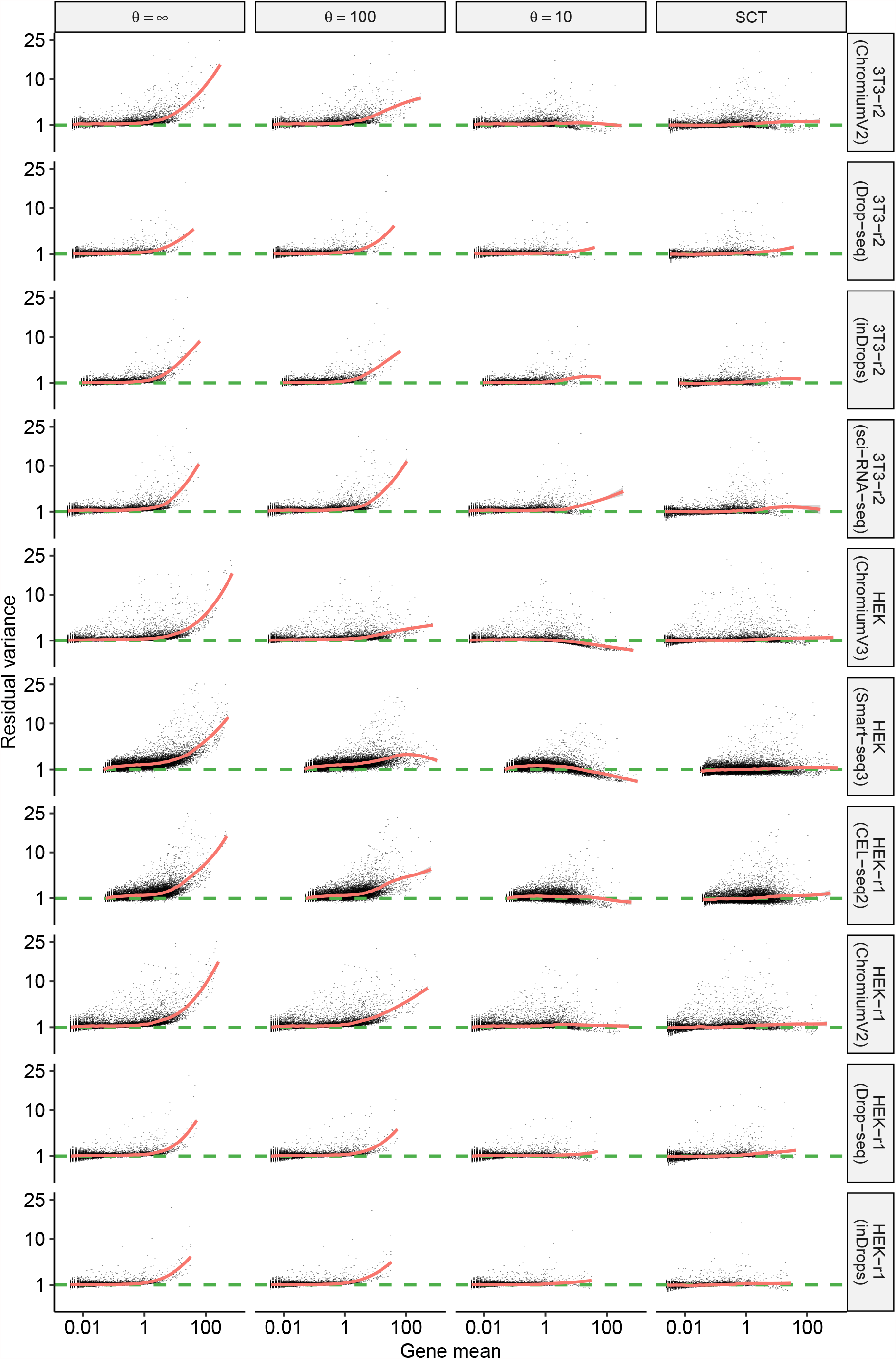
Evaluating variance stabilization for different error models. Same as in Figure 2C, but with additional datasets 11-20. Also shown are results from sctransform (v2 regularization).

**Figure S7.**
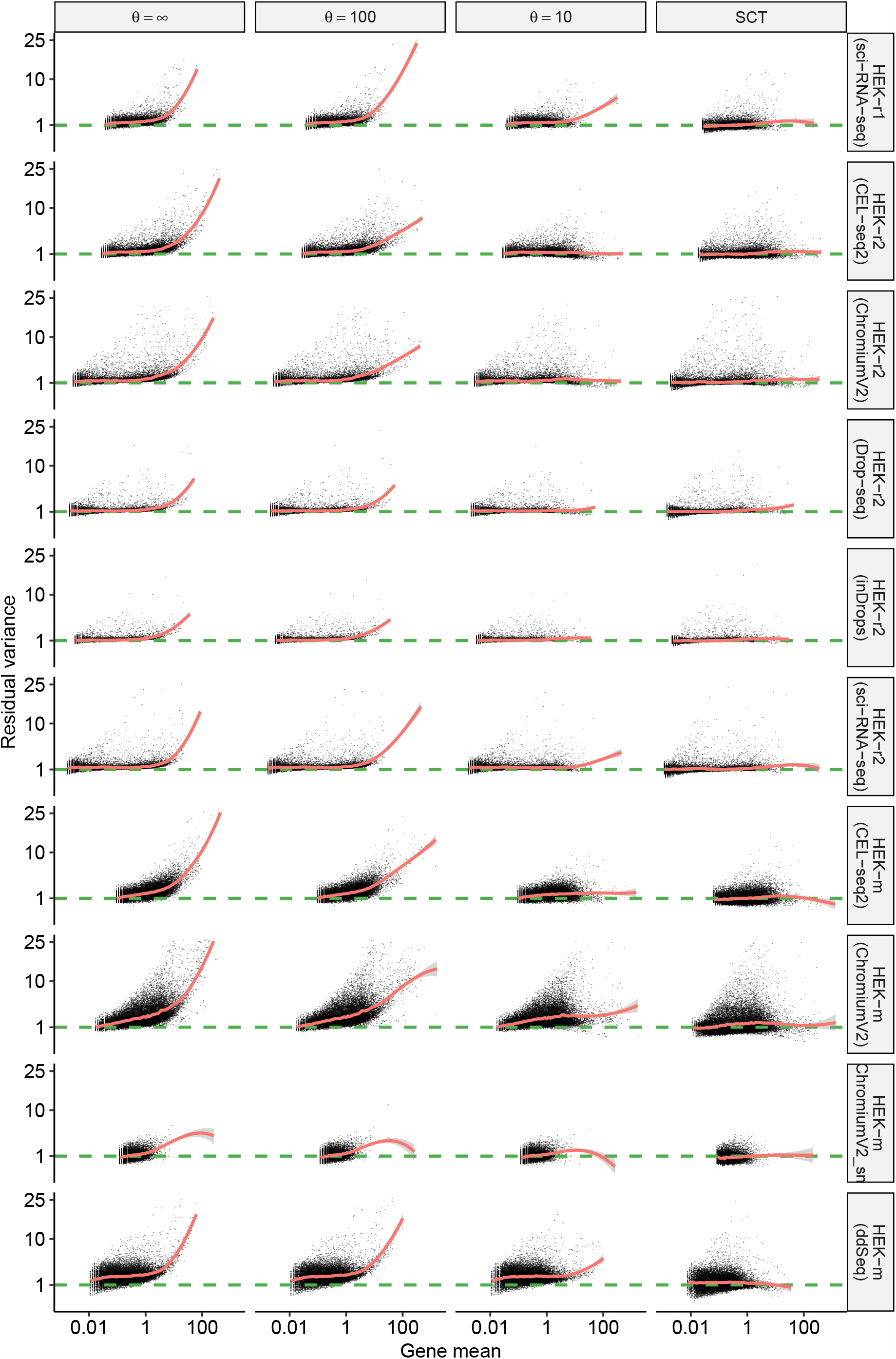
Evaluating variance stabilization for different error models. Same as in Figure 2C, but with additional datasets 21-30. Also shown are results from sctransform (v2 regularization).

**Figure S8.**
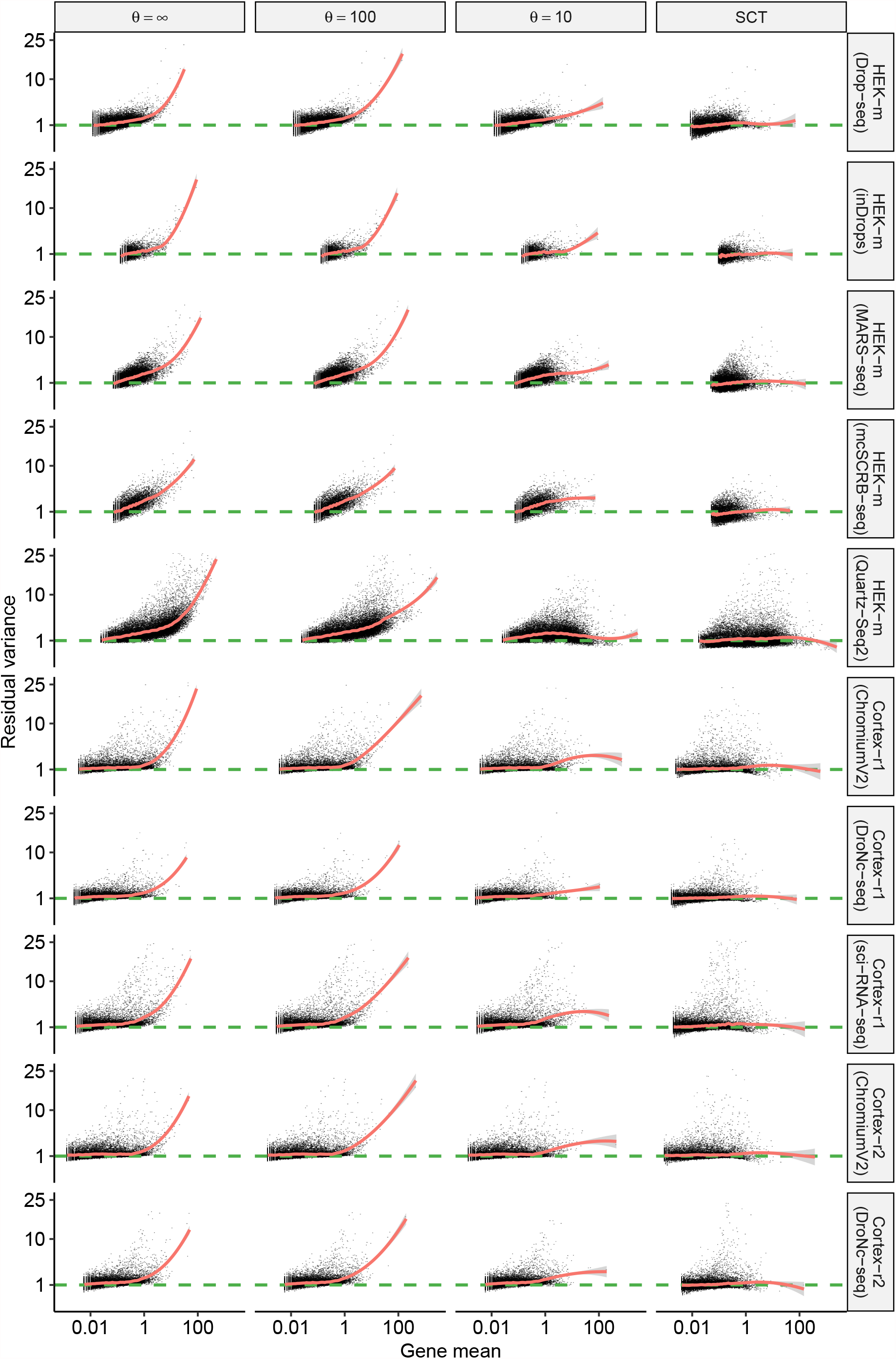
Evaluating variance stabilization for different error models. Same as in Figure 2C, but with additional datasets 31-40. Also shown are results from sctransform (v2 regularization).

**Figure S9.**
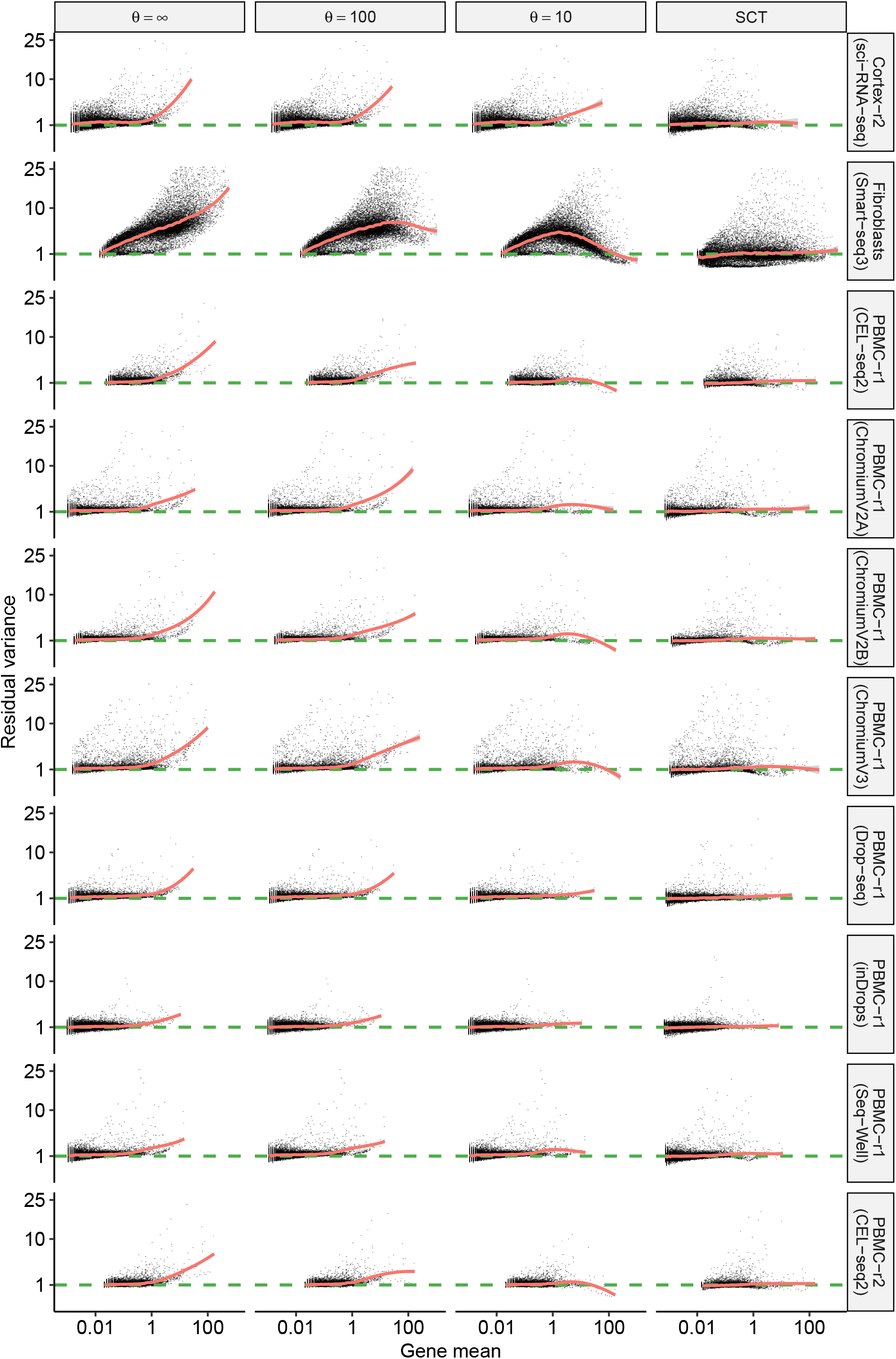
Evaluating variance stabilization for different error models. Same as in Figure 2C, but with additional datasets 41-50. Also shown are results from sctransform (v2 regularization).

**Figure S10.**
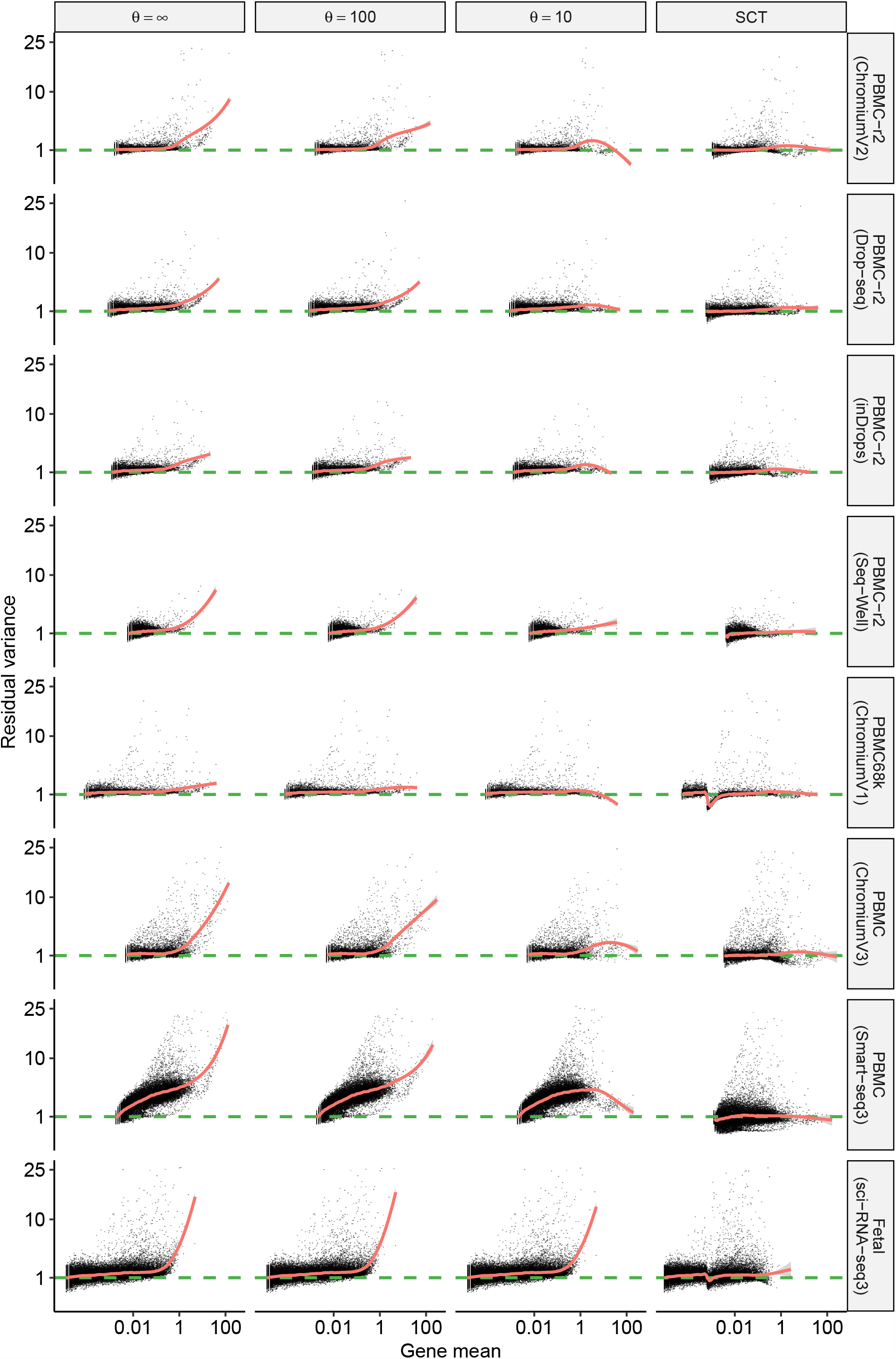
Evaluating variance stabilization for different error models. Same as in Figure 2C, but with additional datasets 51-58. Also shown are results from sctransform (v2 regularization).

**Figure S11.**
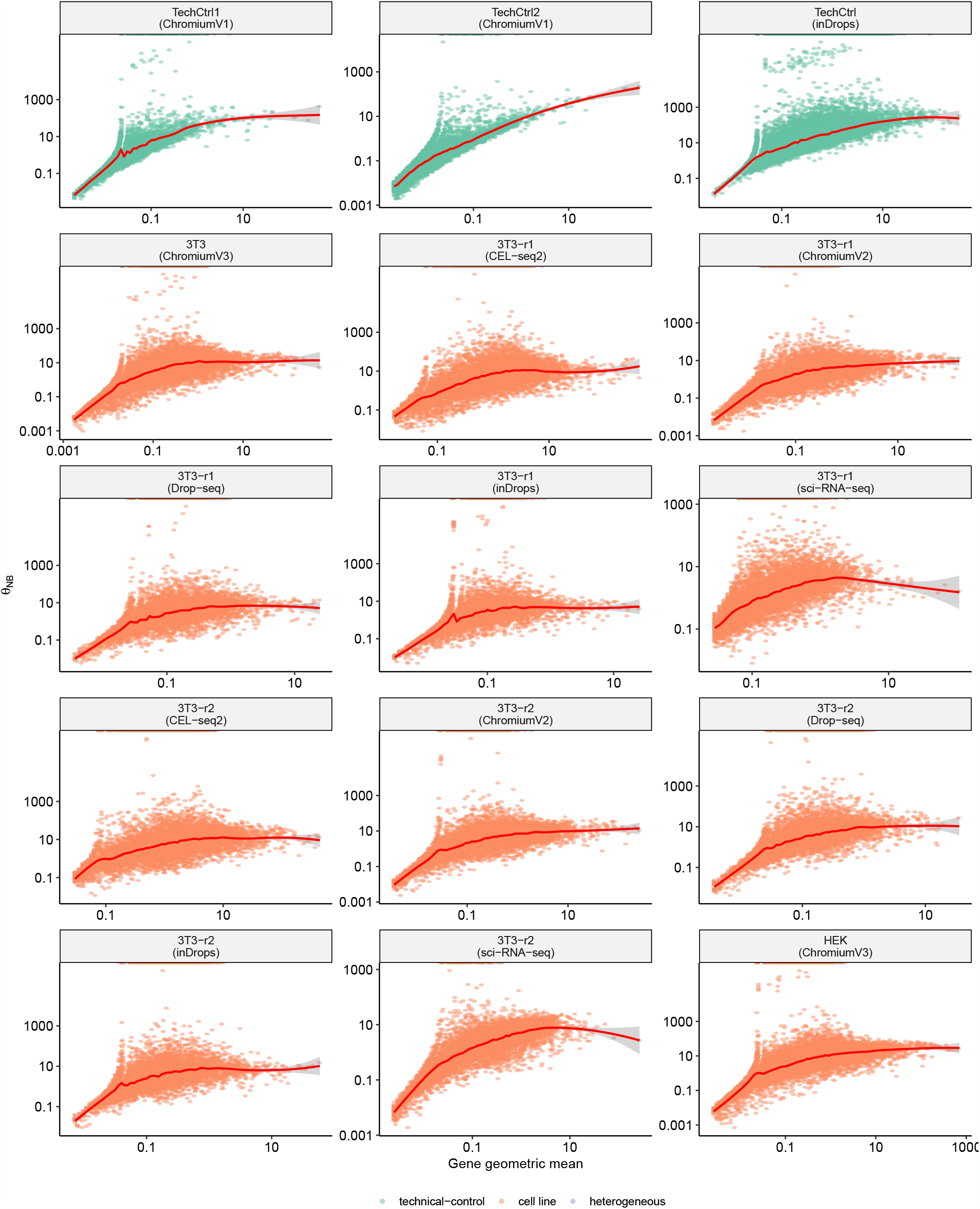
Relationship between inverse overdispersion parameter *θ* and gene abundance *µ*. Overdispersion was estimated using all cells after accounting for library size using a negative-binomial GLM. The red curve indicates a LOESS fit. All datasets exhibit a relationship between gene mean and *θ* [Datasets 1-15].

**Figure S12.**
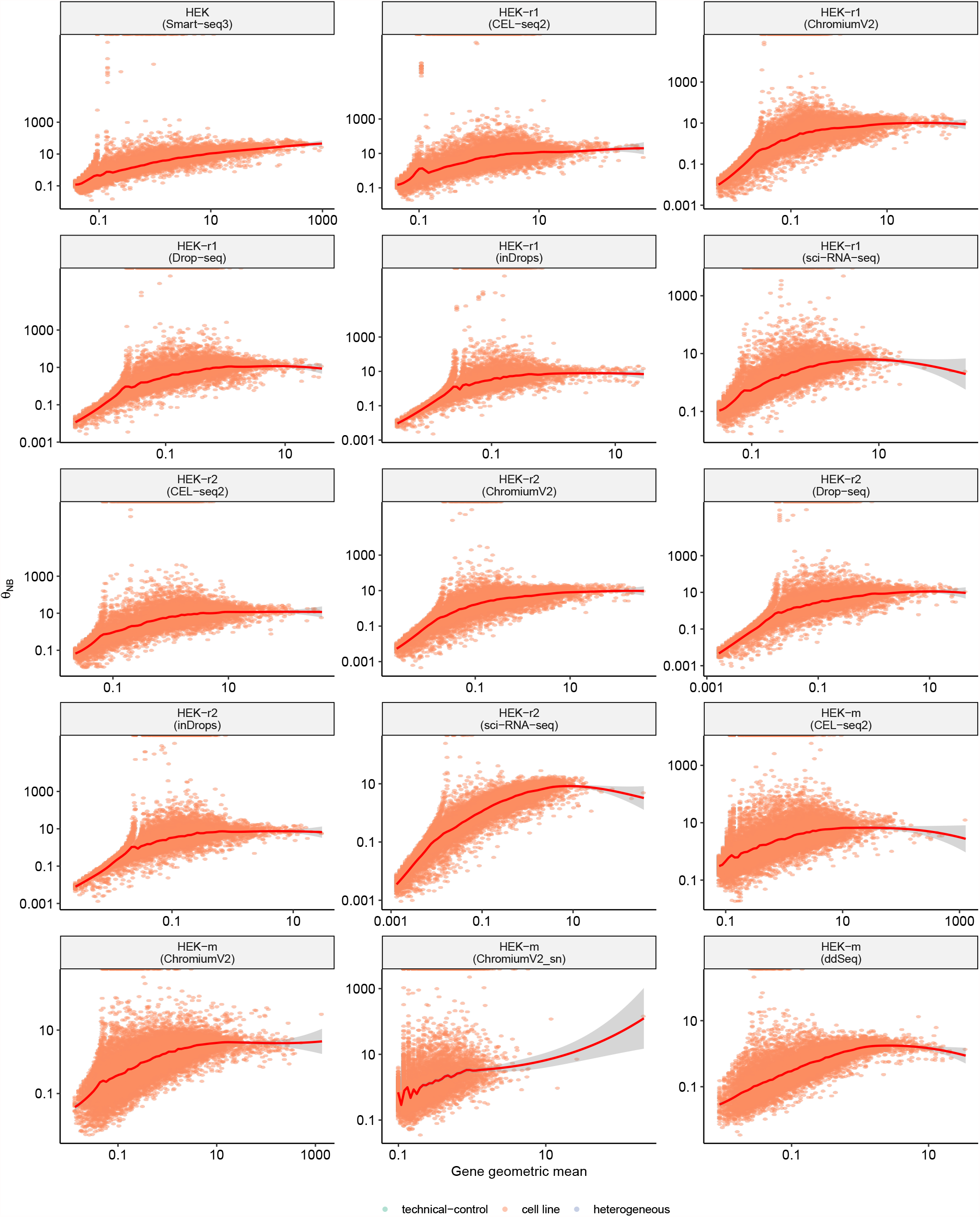
Relationship between inverse overdispersion parameter *θ* and gene abundance *µ*. Overdispersion was estimated using all cells after accounting for library size using a negative-binomial GLM. The red curve indicates a LOESS fit. All datasets exhibit a relationship between gene mean and *θ* [Datasets 16-30].

**Figure S13.**
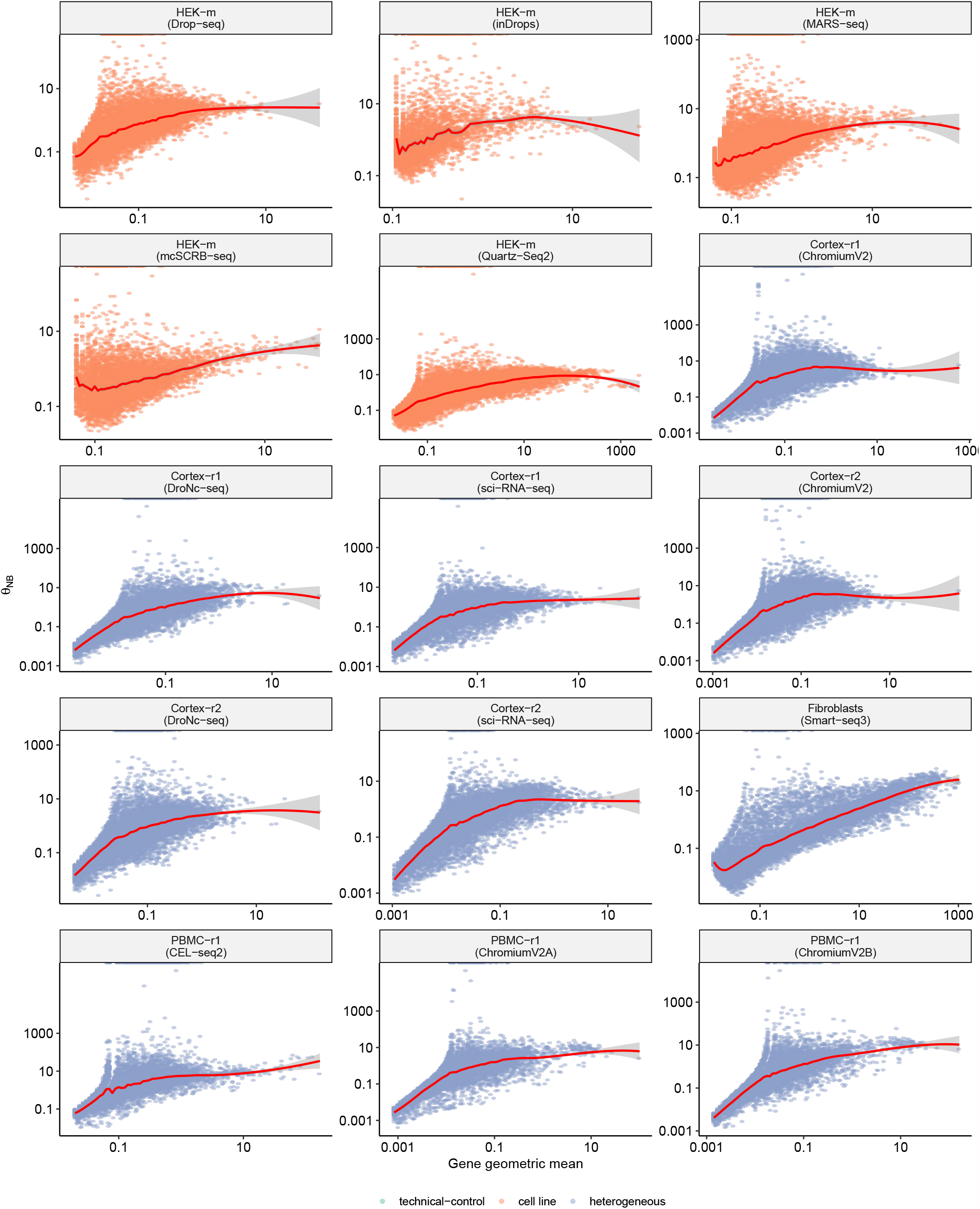
Relationship between inverse overdispersion parameter *θ* and gene abundance *µ*. Overdispersion was estimated using all cells after accounting for library size using a negative-binomial GLM. The red curve indicates a LOESS fit. All datasets exhibit a relationship between gene mean and *θ* [Datasets 31-45].

**Figure S14.**
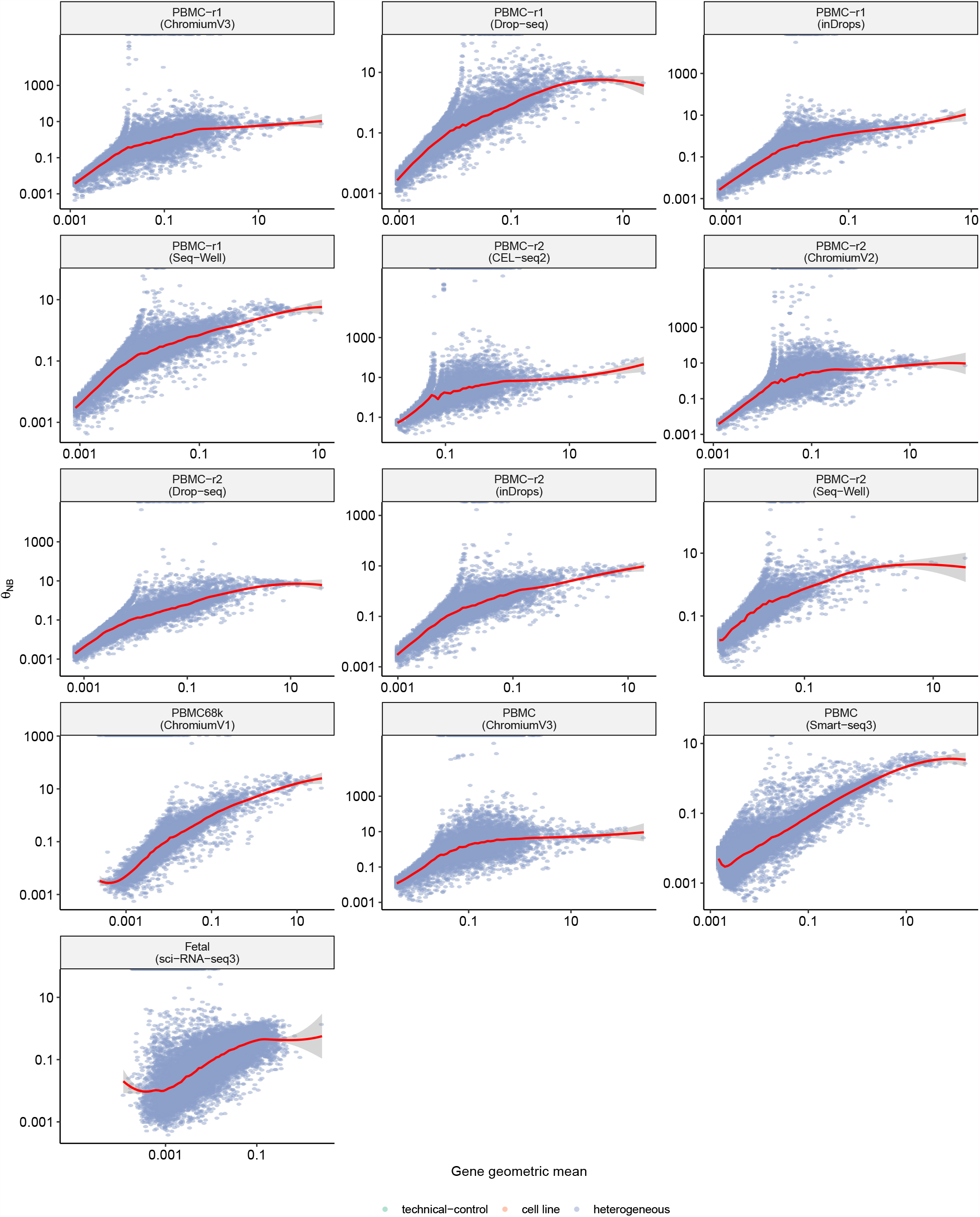
Relationship between inverse overdispersion parameter *θ* and gene abundance *µ*. Overdispersion was estimated using all cells after accounting for library size using a negative-binomial GLM. The red curve indicates a LOESS fit. All datasets exhibit a relationship between gene mean and *θ* [Datasets 46-58].

**Figure S15.**
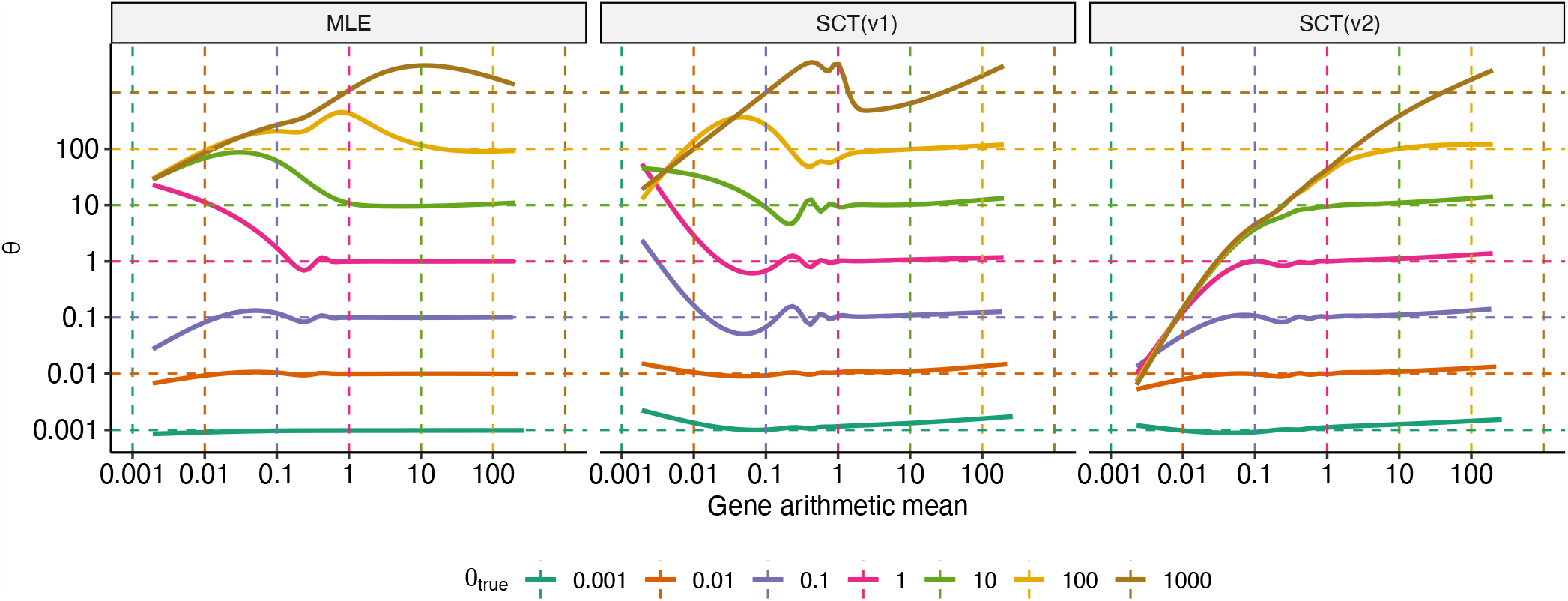
Estimation of dispersion in simulated datasets. Using mean counts distribution from PBMCs profiled using Smart-seq3, synthetic counts matrices were generated using a fixed *θ* = {0.001, 0.01, 0.1, 1, 10, 100}. There is a bias in estimated *θ* from all the three methods: MLE (glmGamPoi (49)), SCT (sctransform) and SCT2 (sctransform, v2 regularization). The bias arises from difficulty in estimating the true *θ* when *µ <* 1 and *µ < θ*. We note that the variance of the NB model is given by 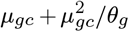. The second term approaches 0 for small values of *µ*, which is where we observe this bias. Therefore, the bias in parameter estimation has minimal impact on both the expected NB variance, and the final Pearson residuals (50).

**Figure S16.**
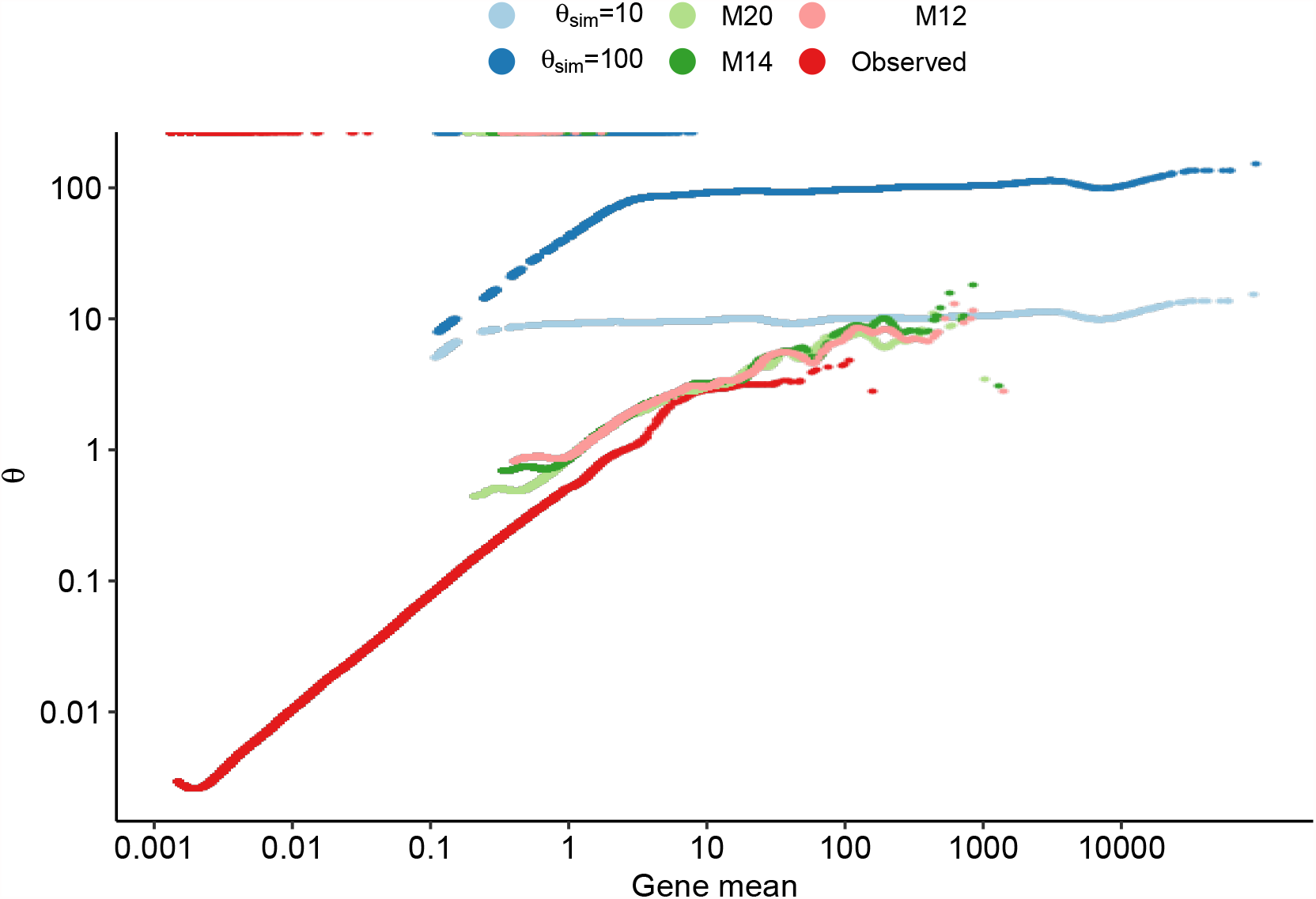
Effect of ‘upsampling’ on *µ − θ* relationship. Relationship between gene mean and dispersion observed in PBMC Smart-seq3 dataset, simulated dataset with different true dispersions (*θ*_sim_ = 10 and 100) and Metacells (M20, M14, M12). The simulated datasets were generated using mean counts from the observed PBMC Smart-seq3 dataset, but by ‘upsampling’ the means to be 500 times larger. Metacells were generated using MetaCell (37) using different parameters of *K* for the KNN graph. M20, M14, and M12 represents 20, 14, and 12 metacells constructed using K = 200, 300, and 400 respectively.

**Figure S17.**
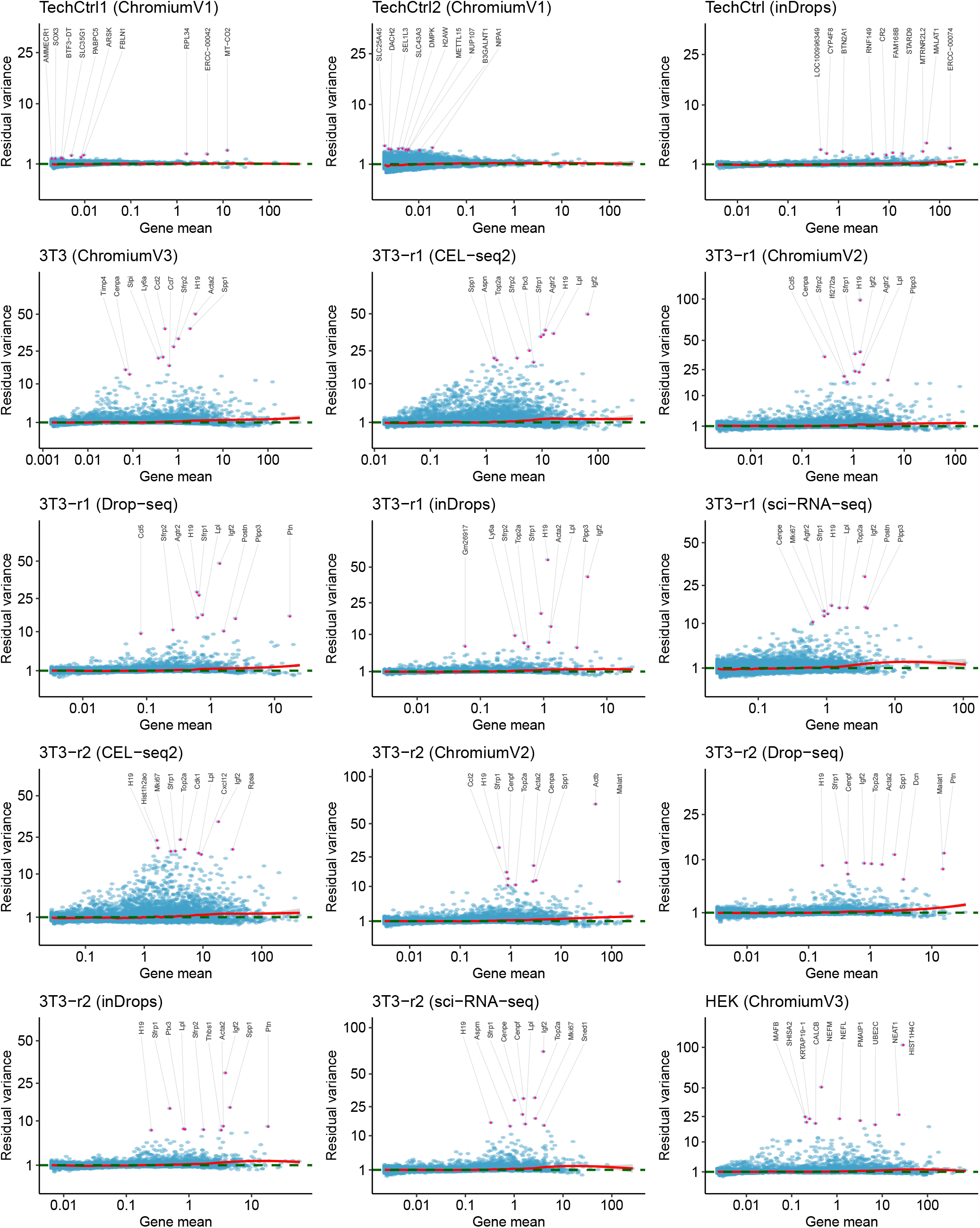
Variance stabilization achieved by sctransform2 across datasets. Y-axis shows variation of pearson residuals as estimated by sctransform (v2 regularization) for datasets 1-15. Top 10 genes with highest residual variances are highlighted.

**Figure S18.**
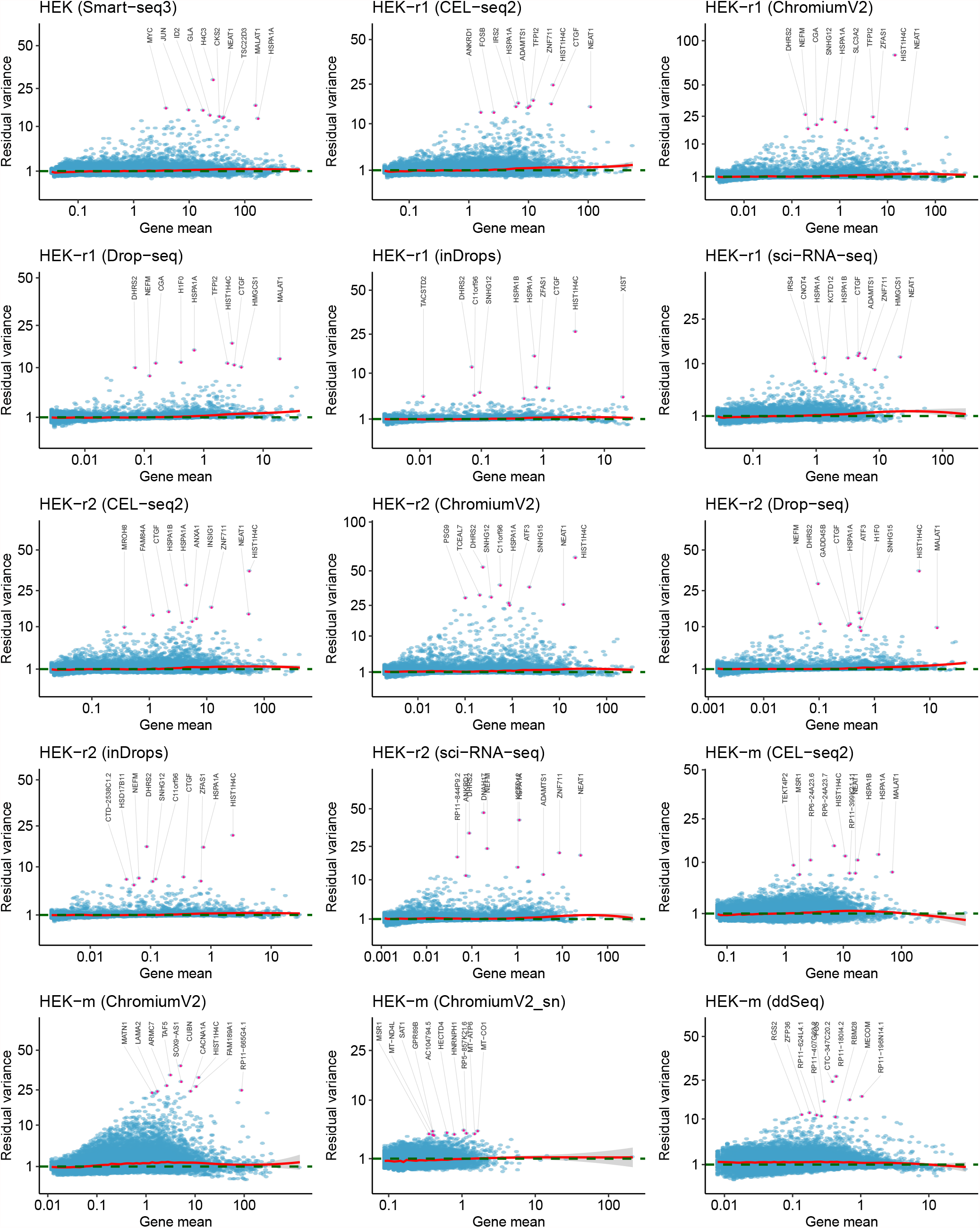
Variance stabilization achieved by sctransform2 across datasets. Y-axis shows variation of pearson residuals as estimated by sctransform (v2 regularization) for datasets 16-30. Top 10 genes with highest residual variances are highlighted.

**Figure S19.**
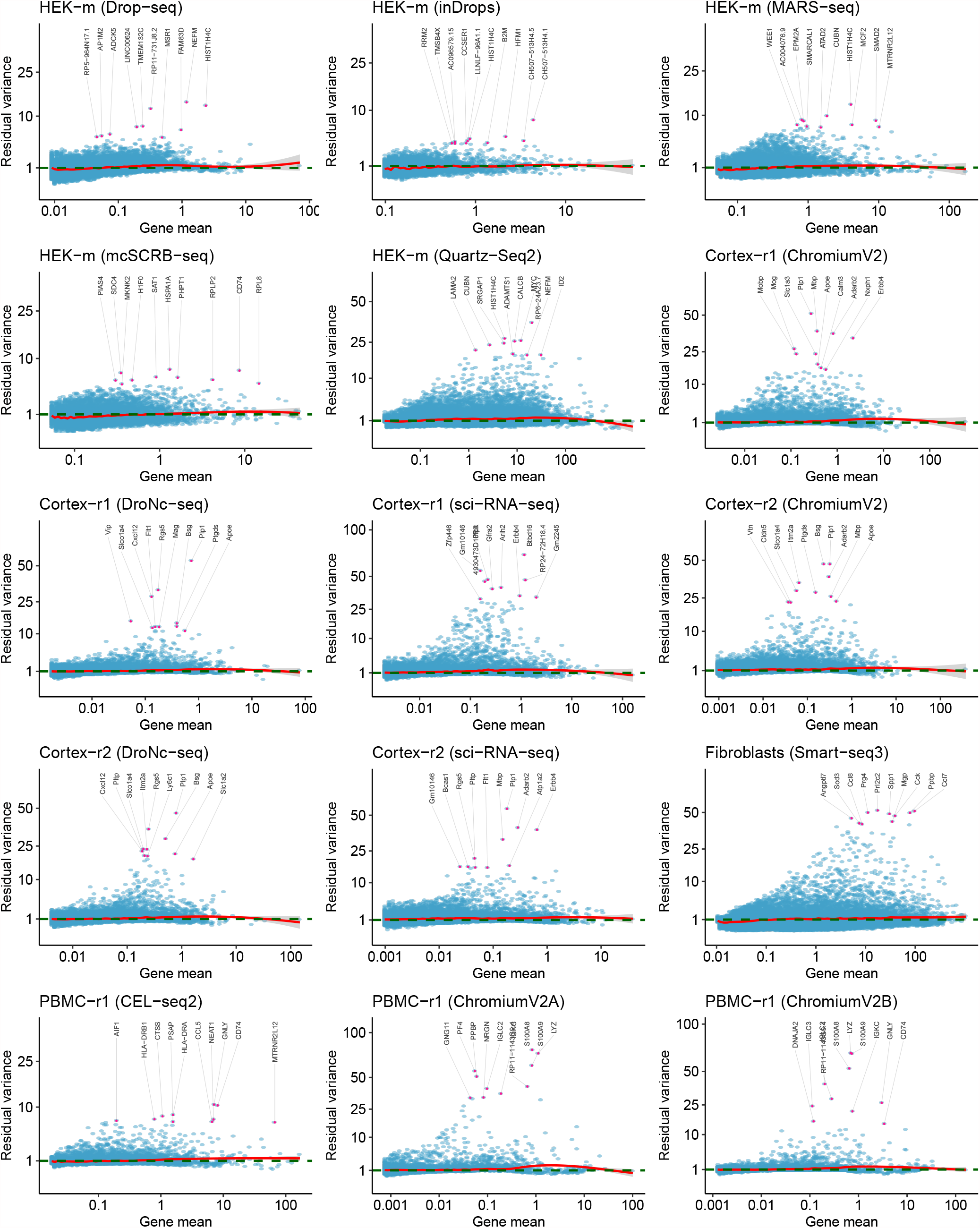
Variance stabilization achieved by sctransform2 across datasets. Y-axis shows variation of pearson residuals as estimated by sctransform (v2 regularization) for datasets 31-45. Top 10 genes with highest residual variances are highlighted.

**Figure S20.**
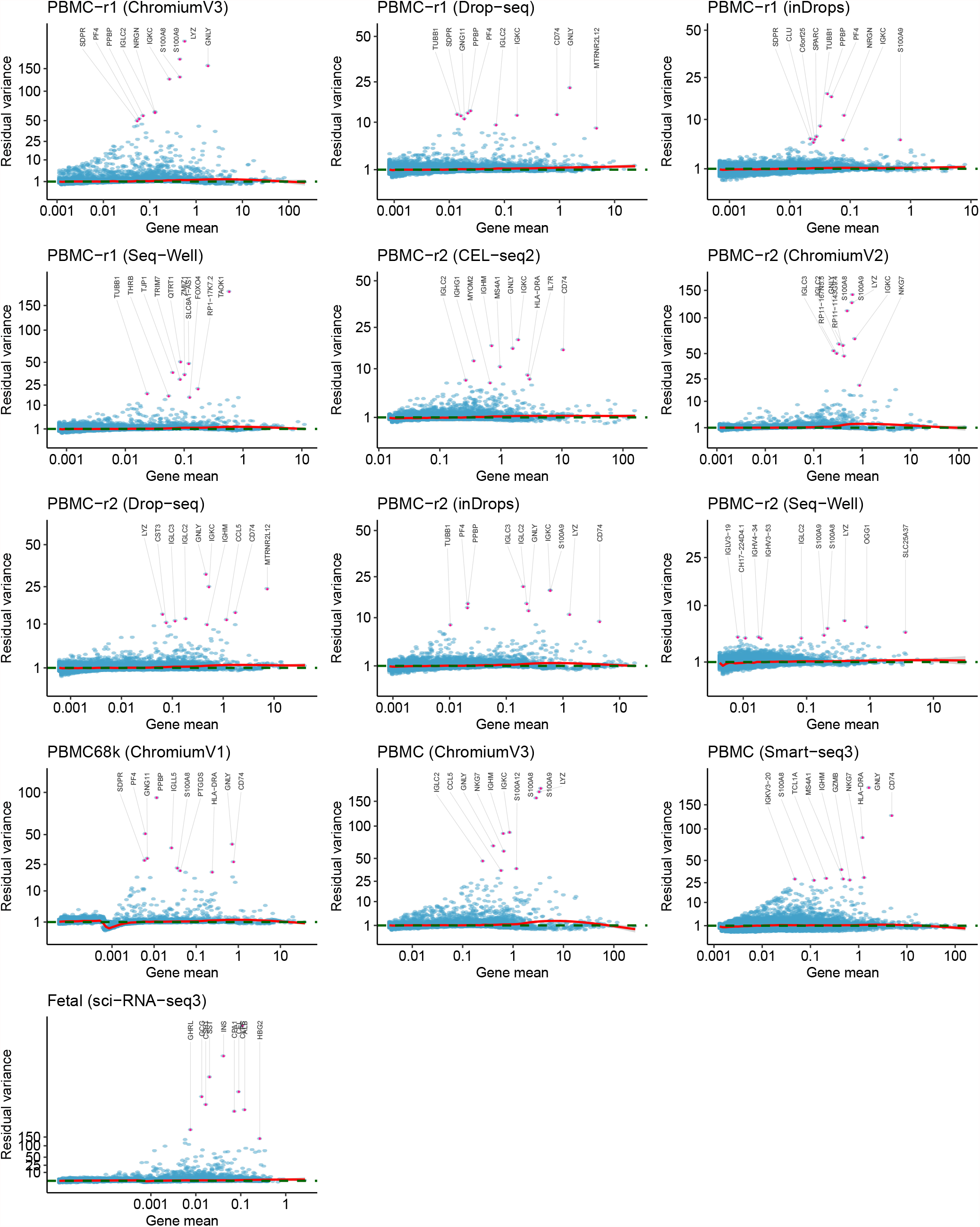
Variance stabilization achieved by sctransform2 across datasets. Y-axis shows variation of pearson residuals as estimated by sctransform (v2 regularization) for datasets 46-58. Top 10 genes with highest residual variances are highlighted.

**Table S1.**
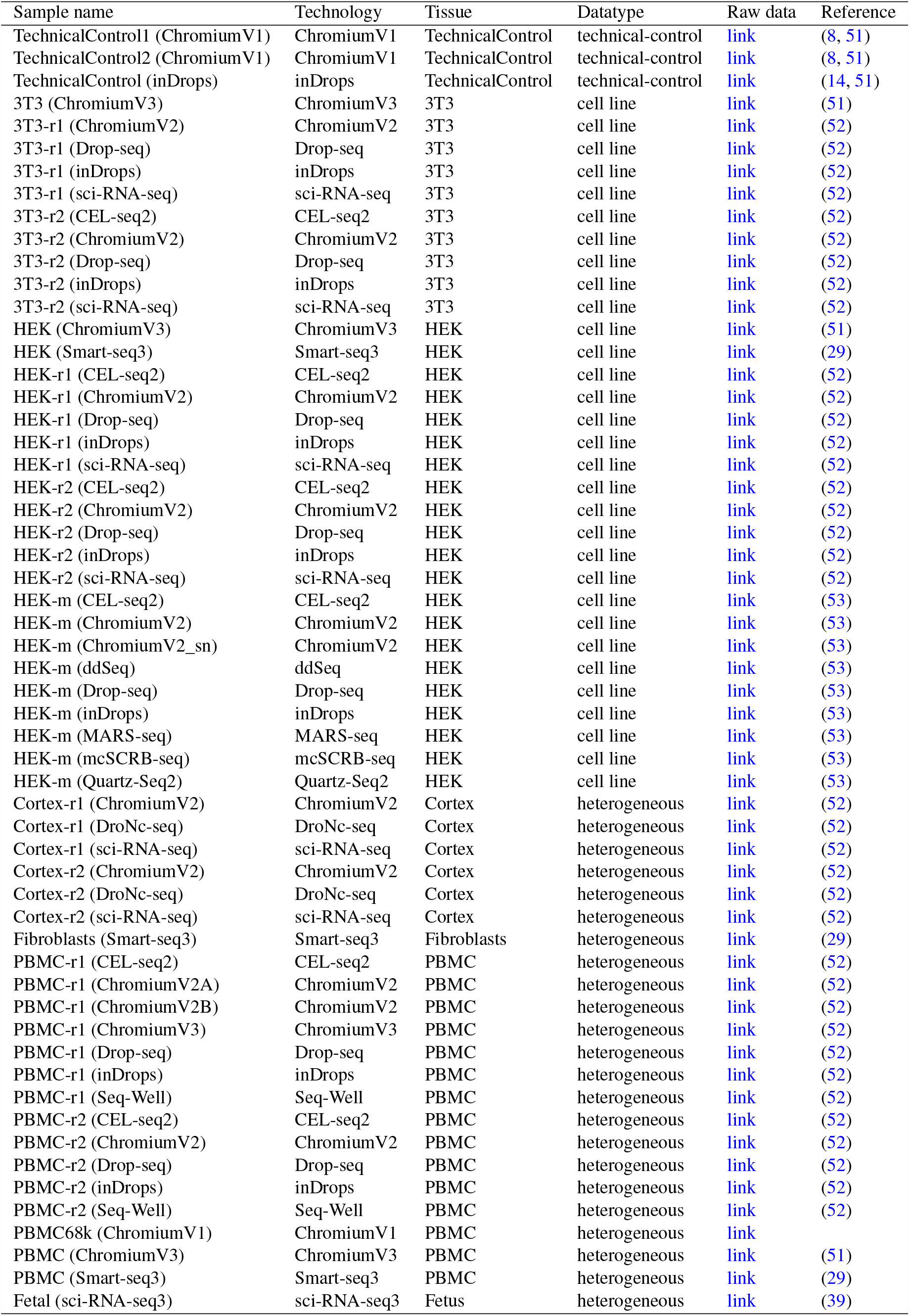
List of datasets used in this study. Raw data can be downloaded from the hyperlinks under ‘raw data’ column. Similar sample names with ‘-r1’ and ‘-r2’ denote replicates from Ding et al.(52) study.

**Table S2.**
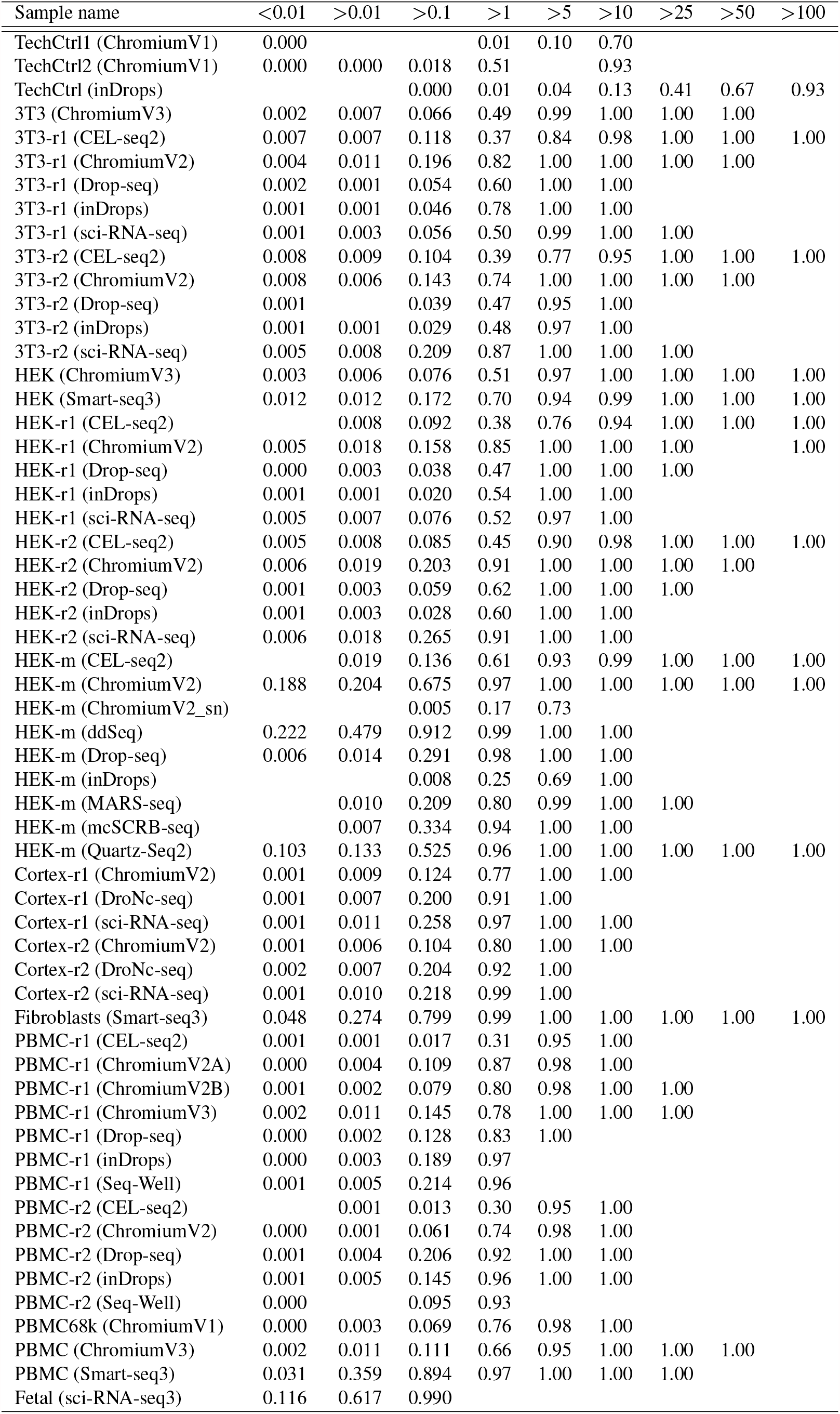
Proportion of non-poisson genes across different gene-mean bins. Columns indicate non-cumulative gene abundance bins between two consecutive labels (for example, *>* 1 refers to all genes with mean *>* 1 and *≤* 5). Each cell entry summarizes the total proportion of genes belonging to a mean abundance bin that were detected to be non-poisson for a dataset.

